# High-throughput discovery of transmembrane helix dimers from human single-pass membrane proteins with TOXGREEN sort-seq

**DOI:** 10.1101/2025.04.22.650048

**Authors:** Samantha M. Anderson, Joshua Choi, Emma M. Cushman, Megan Leander, Srivatsan Raman, Alessandro Senes

## Abstract

The oligomerization of the transmembrane helices of single-pass membrane proteins is crucial to biological function and its misregulation can lead to many diseases. The study of transmembrane helix oligomerization is facilitated by the availability of genetic reporter assays, which are essential tools for understanding the organization and biology of single-pass systems. In particular, reporter assays are crucial for mapping the oligomerization interfaces of transmembrane helices through scanning mutagenesis but their application is limited by the need to clone and measure each construct individually. Here, we present “TOXGREEN sort-seq”, a high-throughput version of the TOXGREEN assay that enables the direct measurement of transmembrane helix oligomerization in large libraries using fluorescence-activated cell sorting and next-generation sequencing. We show that TOXGREEN sort-seq is robust and reproduce the direct measurements of individual constructs with good accuracy and sensitivity. The method produced high-quality mutational profiles from a library of 17,400 constructs designed to probe the interface of 100 potential GAS_right_ dimers predicted from sequences of human single-pass membrane proteins. We report the validated structural model of twelve dimers involved in a variety of biological functions, including immune response (interleukin-22 receptor subunit alpha-1, butyrophilin-like protein 3, hepatitis A virus cellular receptor 2), transport (transferrin receptor protein 1), and cell-surface signaling and proliferation (syndecan-3; semaphorins 5A, 6B and 6D). Remarkably, all three semaphorins in the dataset formed strong dimers and produced mutational profiles consistent with the computational structure. These findings open the possibility that dimerization may be relevant to these proteins’ activity and provide a validated interface for assessing their biological role.

## Introduction

Proteins containing a single transmembrane (TM) helix are the most common type of membrane protein, accounting for over 20% of the total in humans and most organisms^1^. The TM helices of these “single-pass” membrane proteins often play active roles in biological function through the formation of oligomeric complexes^2^. One example is the large number of single-pass receptors whose activity often occurs through ligand-induced stabilization of dimeric conformations^3–5^. Several biophysical techniques have been developed to study the oligomerization of TM helices *in vitro*, using physical methods such as Förster Resonance Energy Transfer^6–13^, sedimentation equilibrium analytical ultracentrifugation^14,15^, disulfide exchange equilibrium^16,17^ and steric trapping^18,19^. Genetic reporter systems are a popular complementary approach to measure the oligomerization propensity of TM helices *in vivo* in bacteria, by linking TM association to gene expression. While less quantitative than the biophysical methods, genetic assays enable faster experimentation since they do not require over-expression, purification, reconstitution, or labeling of proteins. They also have the advantage of measuring TM helix interactions in a natural membrane.

The first genetic reporter assay for measuring TM helix oligomerization was developed by Langosh and colleagues^20^. It was based on the expression in *Escherichia coli* of a fusion protein containing the TM helix of interest connected to the intracellular domain of ToxR, a membrane-bound transcriptional activator from *Vibrio cholera^21^*. In this system, TM helix oligomerization leads to the quantifiable expression of β-galactosidase. Derivatives of the system include versions suitable for characterizing hetero-oligomeric and antiparallel TM helix interactions^22–24^ and a variant in which ToxR can act as either an activator or a repressor by binding to alternative promoter sequences^25,26^. Another ToxR-based system, TOXCAT^27^, expresses an antibiotic resistance gene, enabling the selection of strongly oligomerizing TM helices in large libraries^28–30^. Other variants include systems in which the reporter gene was replaced by luciferase^31^ or Green Fluorescent Protein (GFP)^32^ for improved detection, along with a dominant-negative ToxR system for detecting heterologous association^33^. Other genetic assays use different transcriptional regulators, including GALLEX, which utilizes both a wild-type (WT) and mutated DNA-binding domain of the LexA transcriptional repressor to report heterologous association^34^, and AraTM, which promotes Type I orientation and allows for the assessment of both transmembrane and juxtamembrane regions in oligomerization^35,36^.

These genetic reporter assays are particularly useful for scanning mutagenesis, which is a practical and effective strategy for experimentally mapping the interaction interface of TM helical dimers. Mutagenesis data can validate or inform the development of accurate computational models, which are particularly useful given that single-pass proteins remain severely underrepresented in the structural database^37^. Engelman and coworkers pioneered this strategy using SDS-PAGE as the experimental method for measuring oligomerization. The pattern of disruption in SDS-resistant self-association identified the association interface and supported the computation of early models of the glycophorin A (GpA) dimer^38,39^ and the phospholamban pentamer^40,41^. The development of genetic reporter assays has since greatly streamlined the scanning mutagenesis strategy and it also expanded its applicability to dimers of moderate stability, which are unstable in SDS detergent. These genetic assays also enhanced the phenotype of certain mutants, particularly those of mildly polar residues (G, S, T and Y) that are common in membrane proteins and tolerated in a membrane environment^27^ but can be disruptive irrespective of their position in helices solubilized in a detergent^38,40^.

Just as genetic assays made it easier to map the interface of individual TM oligomers through scanning mutagenesis, the development of high-throughput methods opens the opportunity to extend scanning mutagenesis to large libraries of helices, enabling their simultaneous analysis with comparable effort. TOXCAT is already suited for analyzing large sequence libraries, although indirectly through the use of selection. The expression of the Chloramphenicol AcetylTransferase (CAT) as a reporter gene can be inferred through survival in media containing the antibiotic chloramphenicol, coupled with Next-Generation Sequencing (NGS) to determine the constructs that survive at increasing antibiotic concentrations. Using this strategy, Fleishman and colleagues performed full mutational scanning of three single-pass model systems, exploring the energetics of membrane insertion and dimerization^30^.

Although CAT resistance provides an indirect measure of oligomerization propensity, it would be useful to precisely measure reporter gene expression in high-throughput. Here we present TOXGREEN sort-seq, a high-throughput method that measures reporter gene expression directly. The method is based on the GFP-expressing variant of TOXCAT, TOXGREEN^32^. In traditional TOXGREEN, GFP expression is measured in a fluorescence microplate reader for each construct individually (Fig. 1a). TOXGREEN sort-seq significantly improves the throughput by using Fluorescence-Activated Cell Sorting (FACS) to read all individual cells in a large library and sort them into bins based on their fluorescence. NGS then quantifies the occurrence of each construct within these bins, and the fluorescence of each individual construct in the library is reconstructed using statistical methods.

**Fig. 1.**
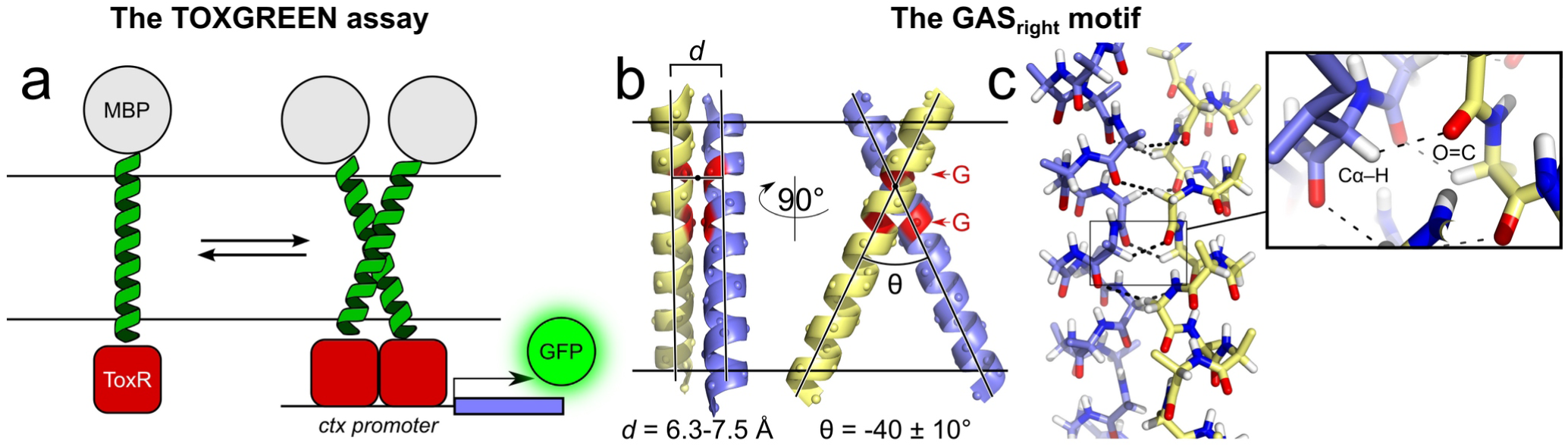
a) The TOXGREEN assay is a derivative of TOXCAT, a widely used genetic assay to study interactions of transmembrane helices within the inner membrane of the bacterium *Escherichia coli*. The assay is based on a fusion construct that links a transmembrane domain of interest with a cytoplasmic DNA-binding domain from the Vibrio cholerae ToxR protein. Interaction driven by the transmembrane domain results in dimerization of the ToxR domain, which, in turn, activates the expression of the reporter gene superfolded GFP. The resulting fluorescence is used as a measure of the ability of the transmembrane domain to self-associate.

We applied TOXGREEN sort-seq to validate the association interface of 100 computationally predicted human TM helix dimers in high-throughput using scanning mutagenesis. The opportunity for a large-scale validation of structural models of TM dimers was provided by a large dataset of potential TM dimers that were previously predicted from the sequences of all single-pass proteins in the human proteome using the program CATM^42,43^. These dimers were predicted to associate through the formation of GAS_right_ structures, a common dimerization motif for single-pass proteins^44,45^ illustrated in Fig. 1b. GAS_right_ dimers are characterized by a right-handed interaction angle and signature GxxxG and GxxxG-like sequence motifs (GxxxG, GxxxA, SxxxG, etc.)^46–48^. Their helical backbones are in close contact and the interactions are stabilized by van der Waals packing and networks of inter-helical weak hydrogen bonds between Cα–H carbon donors and carbonyl acceptors (Cα–H···O=C) (Fig. 1c)^42,43^. The strength of association of GAS_right_ dimers is modulated by its geometry and sequence^42^, forming both strong structural dimers^49,50^ as well as weak dynamic systems^4,51–56^. The dataset contains a range of dimers with different estimated energy scores, allowing for testing the scanning mutagenesis approach across a range of association strengths. Further, the selected proteins include many that are biologically and biomedically relevant, but are not yet known to form dimers or if they dimerize specifically through their TM helices.

Our analysis demonstrates that the statistically reconstructed fluorescence signals obtained with TOXGREEN sort-seq reproduce the direct fluorescence measurements of individual constructs with good accuracy and sensitivity. We show that the method is effective for measuring thousands of constructs consisting of WT sequences and point mutations simultaneously, generating high-quality, reproducible mutational profiles. Finally, we present the structural model of twelve GAS_right_ dimers of previously unknown structure that have been validated by our mutational analysis. These proteins participate in a variety of important processes, including transport, the modulation of immune response and cell-surface signaling, proliferation and adhesion. In particular, we discover that the TM helix of three different semaphorins is a strong dimerization domain. Semaphorins are a family of cellular surface proteins that mediate cell-to-cell communication and are critical for regulating cell morphology, cell differentiation and cell-cell interactions both in neuronal and non-neuronal cell types. They are known to dimerize through interactions in their extracellular domains. Our data indicates that the TM domain of these semaphorins is an additional dimerization domain and suggests it may be important for the assembly and signaling of these proteins.

## Results and Discussion

### The TOXGREEN sort-seq method

The TOXGREEN sort-seq method is schematically illustrated in Fig. 2. The experiment starts with a synthetic DNA oligo pool library in which each oligonucleotide encodes a unique TM helix sequence (although the method is applicable to other approaches for generating libraries, such as randomized mutagenesis). The pool of DNA inserts is cloned into the standard TOXGREEEN plasmid (pccGFPKan) and transformed into MM39 *E. coli* cells, resulting in a cell library expressing each oligo pool sequence. The cells are then sorted into multiple bins using FACS based on the expression levels of GFP. The DNA for each of these sorted populations is then extracted and analyzed by NGS to obtain a count for each library construct within each bin. As described in Methods, the distribution of the counts of each construct in the bins is then used to mathematically reconstruct their estimated GFP fluorescence, using a statistical inference approach based on a weighted average adapted from Kosuri et al.^57^.

**Fig. 2.**
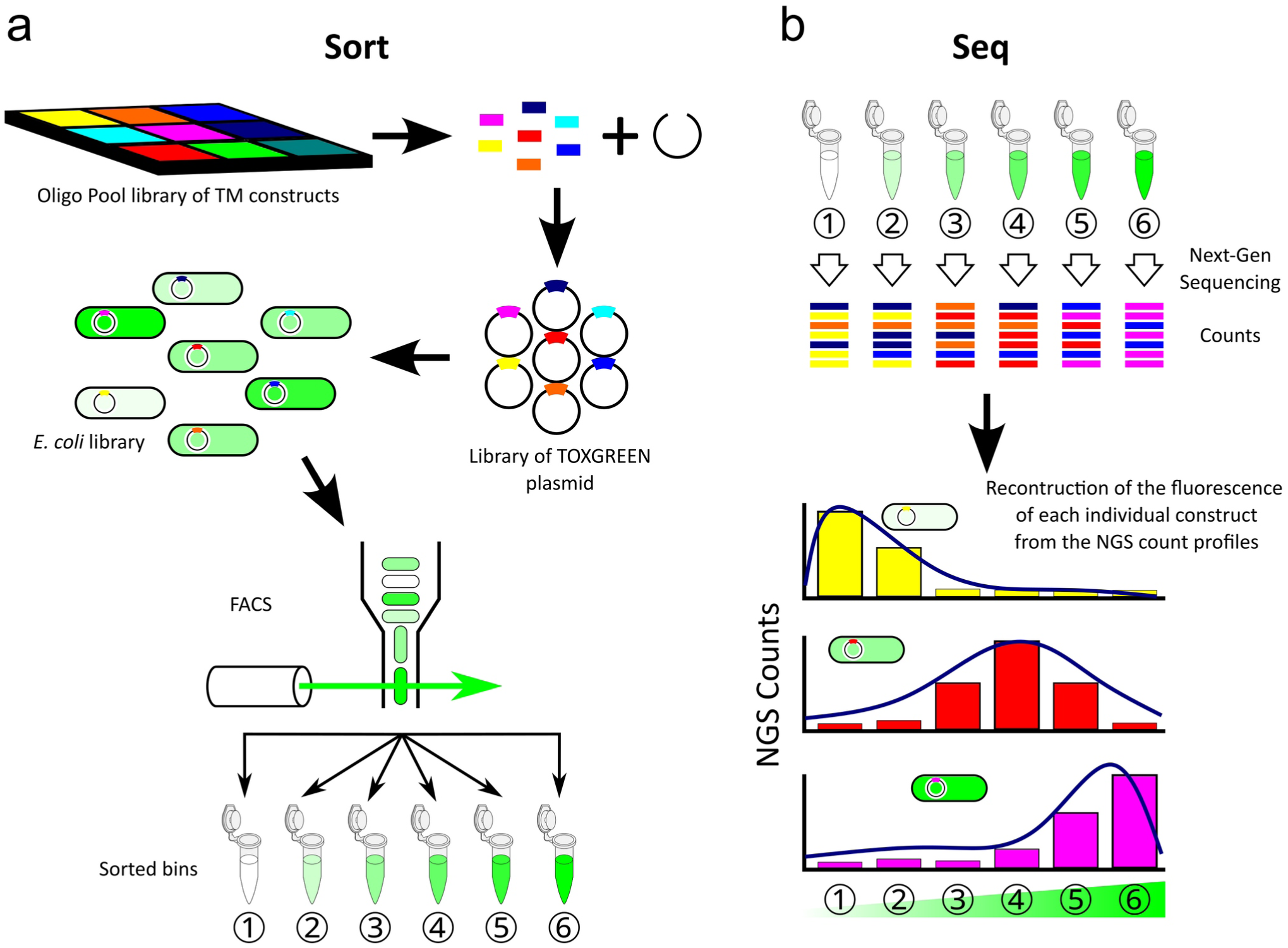
Schematic representation of the TOXGREEN sort-seq method. a) The procedure starts with a library of oligonucleotides encoding the TM helices of interest. These are cloned into a TOXGREEN plasmid and transformed into *E. coli*. The cells are sorted into bins based on their GFP expression using FACS. b) The population of each construct in the bins is determined using Next-Generation Sequencing. The count distribution of each construct in each bin is used to reconstruct the fluorescence of each individual construct.

### High-throughput data accurately recapitulates individual measurements

To assess the performance of TOXGREEN sort-seq, we performed three comparisons of directly measured fluorescence vs its reconstructed values across conditions of increased complexity. For this step, we used a small library of 100 TM helices (called the validation library, supplementary Table S1) that had been individually characterized and displayed varying degrees of GFP expression.

First, to establish the feasibility of using a cell sorter for measuring GFP expression in our systems, we compared the median fluorescence distribution of individual cells measured through flow cytometry with the fluorescence measured traditionally in bulk using a plate reader. As illustrated in Fig. 3a, the comparison resulted in a strong linear correlation between the two instrument measurements (Deming regression, *ρ* = 0.886) with standard deviations (error bars) of similar relative magnitude. The experiment demonstrates that a fluorescence-activated cell sorter has the dynamic range and sensitivity to accurately measure the GFP fluorescence of TOXGREEN constructs.

**Fig. 3.**
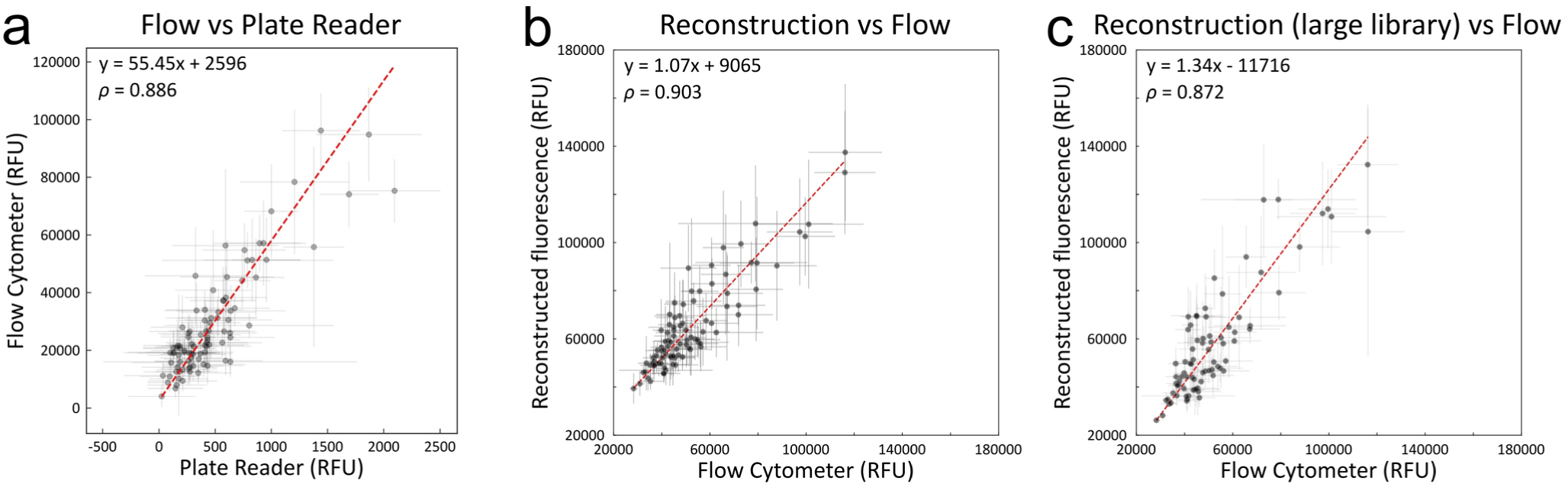
Validation of the TOXGREEN sort-seq method. Validation of the TOXGREEN sort-seq method. A small library of 100 pre-characterized constructs (the validation set) was created for testing the method (Table S1). a) Comparison between direct measurement of GFP fluorescence in a plate reader vs direct measurement of individual constructs in a flow cytometer. b) Direct measurement in flow cytometer vs statistical reconstruction using sort-seq. c) Same as panel b, except that the validation library was mixed with a large library of over 17,000 unrelated constructs. The observed linear relationships establishes that TOXGREEN sort-seq can reconstruct the fluorescence of individual constructs with good accuracy and sensitivity in large pools of sequences. RFU = Relative Fluorescence Units. *ρ* = Pearson correlation coefficient. Linear fits performed using Deming regression. Error bars represent standard deviation of three replicates for the plate reader values and at least five replicates for the flow cytometer values and for the reconstructed fluorescence values.

Next, we assessed TOXGREEN sort-seq’s statistical approach by comparing the fluorescence reconstructed from NGS data with the median fluorescence of each construct directly measured in a cell sorter. We first reconstructed the GFP fluorescence on a small scale by sorting a library containing exclusively the 100 members of the validation library (Fig. 3b). We then proceeded to reconstruct the fluorescence of the validation library when these were mixed with a large library of 17,305 additional sequences (Fig. 3c). Both measurements of the validation set resulted in excellent linear relationships between reconstructed and measured fluorescence (*ρ* = 0.903 and 0.872, respectively). These experiments establish that the TOXGREEN sort-seq method can reliably evaluate the association of TM helices on a massive scale.

### High-throughput evaluation of proper membrane insertion (maltose growth assay)

To assess whether a TOXGREEN construct is expressed in the membrane and inserted with the correct topology, it is standard practice to perform a separate MalE complementation assay^27^. The MM39 *E. coli* strain used in TOXGREEN is Maltose Binding Protein (MBP)-deficient; it can grow in a M9 minimal media where maltose is the only available carbon source only if the chimeric TOXGREEN constructs are expressed and inserted into the inner membrane with the C-terminal MBP moiety oriented to the periplasmic side. Canonically, this assay is performed using M9 maltose selection plates with individually streaked samples, but this approach is not suitable for large libraries. For compatibility, we scaled up the assay by implementing a liquid culture variant where the entirety of the library is grown in liquid M9 maltose minimal media. The make-up of the surviving library members is identified at time points via next-generation sequencing until the non-inserting constructs are out-competed from the population due to poor uptake of the maltose, leading to a massive decrease in enrichment score.

To assess the feasibility, we compared the growth of the constructs in the validation library with the traditional solid M9 media and the new liquid media protocol. We found the same samples that failed to grow on the selection plates (supplementary Fig. S1a) also displayed greatly diminished presence in the population after a period of extended growth in liquid M9 maltose minimal media (supplementary Fig. S1b and c), suggesting combined library growth in liquid media recapitulates the original test. However, we found the necessary growth time and enrichment score to somewhat vary between library samples. Therefore, for calibration with different libraries and populations, we suggest the inclusion of a set of standard non-inserting constructs (such as 1E01, 2E11, 2H11, 2F12 and 1E11, listed in supplementary Table S1) to identify the most appropriate threshold enrichment score. Any samples with similar growth levels to these negative controls in M9 minimal media should be considered as likely poorly expressing or inserting.

### High-throughput mutagenesis study with TOXGREEN sort-seq

We applied TOXGREEN sort-seq to a large-scale scanning mutagenesis analysis aiming to validate the helix-helix interaction interfaces of 100 computationally predicted GAS_right_ TM dimers^42,43^. These potential dimers were selected from a list of 1,141 GAS_right_ interfaces that we previously predicted from all single-pass proteins annotated in the human genome^42^. The 100 TM sequences were selected using the following criteria: i) they had a predicted TM helix of 21 amino acids (the most common length annotated in Uniprot, for consistency); ii) they did not contain more than one strongly polar residue (D, E, H, K, N, Q); iii) they did not contain a proline residue; and iv) their model did not display significant inter-helical interactions in the juxtamembrane polar region. From the remaining sequences, we selected the top 100 (listed in supplementary Table S2) for the mutational analysis based on their CATM energy score.

A wide variety of mutational strategies have been applied in the literature to oligomerizing systems TM helices. Some studies have used very large sets of mutations, up to thirteen^38^ and even seventeen^58^ changes at each position. A TOXCAT study identified the dimerization interface of BNIP3 using eight mutations per position^59^. Our group has typically substituted seven hydrophobic amino acids at all positions (A, I, F, G, L, M and V)^60,61^, but others have used more minimalistic approaches. For example, TOXCAT was used with a combination of alanine- and cysteine-scanning (the latter in combination with cross-linking) to identify the interfaces for the thrombopoietin receptor TM^62^. Another study used either Ala or Ile at each position (depending on size and polarity) to identify the interface of the human B-cell receptor in GALLEX^63^. Others have used scanning mutagenesis with a single amino acid, such as Ala^64^ or Phe^65^. Aiming for an optimally sized library, we decided a middle-ground approach, introducing three mutations at each position, namely to Ala, Ile or Leu. If the WT identity at a position consisted of one of these three amino acids, we introduced an additional Phe mutation to maintain the total of three. As explained below, we also explored the effect of TM insert length in scanning mutagenesis. For this reason, each of the 100 constructs was assessed at its original 21 amino acid length (called the “21-AA” library) and with truncations to 19 (“19-AA”) and 17 amino acids (“17-AA”). The final combination of the number of constructs, the helix sizes and the mutations yields a theoretical library of 17,400 constructs, consisting of 300 WT sequences (100 constructs each tested at three lengths) and 17,100 point mutants.

### Comparison of TOXGREEN signal as a function of construct length

The length and hydrophobicity of the TM helix insert can influence the reporter output in TOXCAT/TOXGREEN^66^, which is critical for sensitivity in scanning mutagenesis. To systematically evaluate how the length of the TM domain affects this outcome, we compared the reconstructed fluorescence output of the same constructs (including WT sequences and all point mutants) in the three libraries. We found that the longer 21-AA constructs displayed significantly higher reconstructed fluorescence (78,000 ± 32,000 RFU) compared to the 19-AA (58,000±22,000 RFU) and 17-AA libraries (52,000±26,000 RFU) (Fig. S2a). Linear regression analysis of the samples across the three lengths (Fig. S3) shows significant proportionality in the fluorescence, indicating that overall there is still correspondence in the relative TOXGREEN output of a sequence at varying lengths. It should be noted, however, that many of the constructs that display moderate to high signals in the 21-AA library are near the baseline when shortened to 17 residues (Fig. S3c). This data suggests that for improved fluorescence output, the 21-amino acid TM helix is preferable. However, as noted below, the analysis of the mutagenesis profiles indicates that the longer length may not always be the optimal choice.

A summary of the mutagenesis analysis is provided in Table S3. The WT TOXGREEN signal of 35 constructs had a reconstructed fluorescence value close to the monomeric baseline in all three lengths and thus these constructs were excluded from the mutagenesis analysis. The mutagenesis profiles of the 65 remaining constructs are reported in supplementary Fig. S4, where phenotypes are classified according to five levels of disruptions, ranging from WT-like (white) to completely disruptive (red). For each position, we calculated a position-specific “disruption score” by averaging the disruption levels of all mutants of that residue. As illustrated in Fig. 4 we identified five types of profiles. Twelve constructs displayed a profile that is consistent with the predicted interface of their GAS_right_ model (Fig. 4a). A comparable number (13) displayed a pattern of disruption that maps on one face of the helix but does not match the predicted GAS_right_ model (Fig. 4b), suggesting that these helices may possibly oligomerize through a different structure. Interestingly, three of these profiles match reasonably well with their AlphaFold 3 (AF3) prediction, suggesting that they are likely to highlight biological interfaces (P78380: Oxidized low-density lipoprotein receptor 1, P16410: Cytotoxic T-lymphocyte protein 4, and Q9ULG6: Cell cycle progression protein 1, illustrated in supplementary Fig. S5). Six constructs had a pattern of disruption that does not map to a clear interface (Fig. 4c). Three constructs displayed a single polar amino acid as the only stringent requirement for oligomerization (Fig. 4d), consistent with the tendency of these amino acids to drive promiscuous association of TM helices^15,67^. Surprisingly, the remaining 31 constructs displayed a minimal level of disruption across all positions (Fig. 4e).

**Fig. 4.**
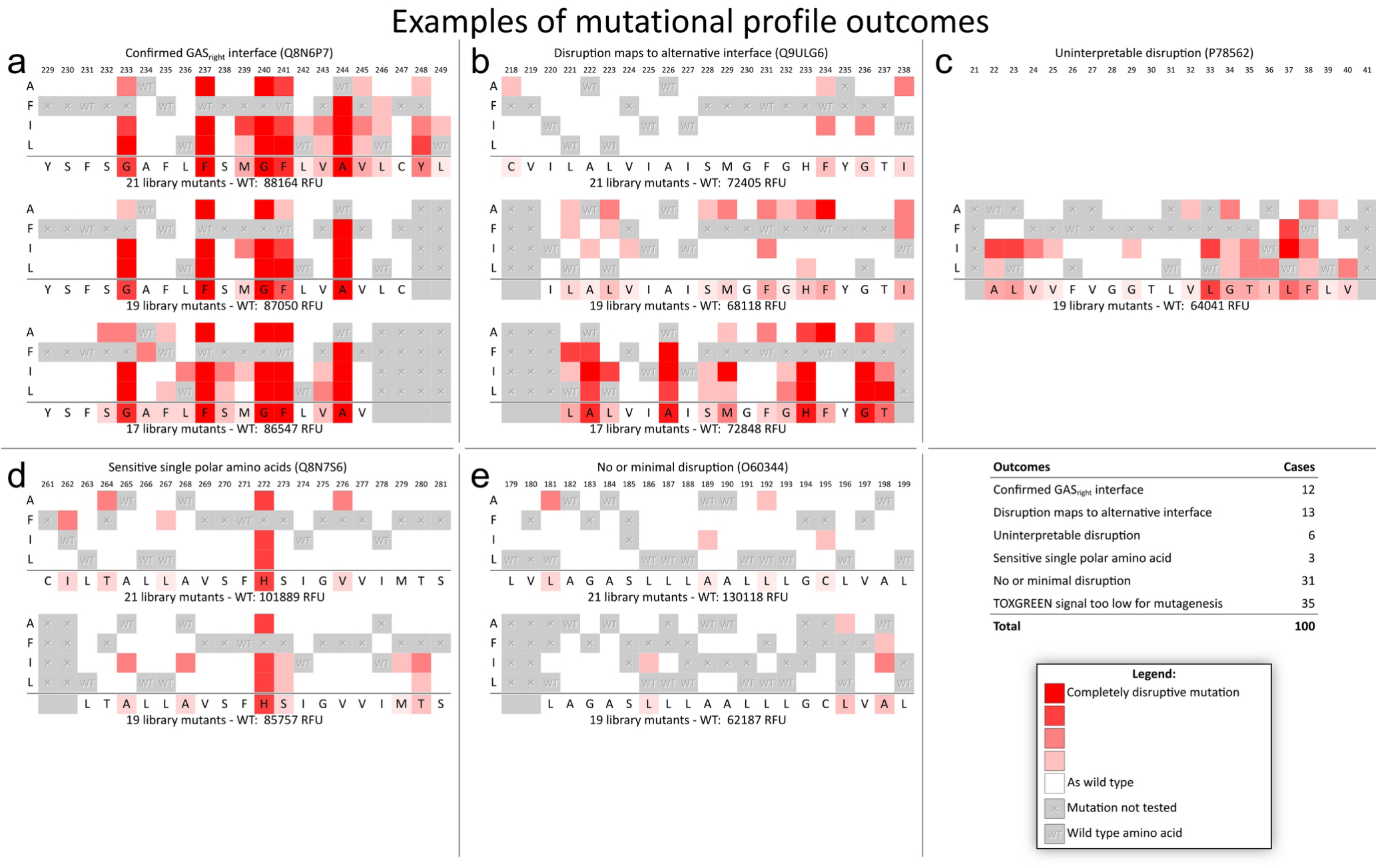
Examples of mutagenesis outcomes. The mutagenesis analysis of 65 constructs produced outcomes that can be classified into 5 groups. a) Twelve constructs displayed a profile that is consistent with the predicted interface of their GAS_right_ model. b) Thirteen constructs displayed a pattern of disruption that maps on one face of the helix but does not match the predicted GAS_right_ model. c) Six constructs had a pattern of disruption that does not map to a clear interface. d) Three constructs displayed a pattern consisting of a single polar amino acid as the only critical position for oligomerization. e) 31 constructs displayed no disruption or a minimal level of disruption across all positions in all the available mutagenesis profiles at various lengths.

When the mutational profiles at different TM lengths are compared, there is an overall notable tendency for mutations to have a stronger effect in the shorter constructs, which is useful for identifying critical positions. The mutational profile of the 21-AA constructs displayed minimal or no disruption in 60% of the cases, whereas it occurs only about 13% and 14% of the 19-AA and 17-AA constructs, respectively. This is evident in the examples in Fig. 4a and b, in which the mutations of the 21-AA constructs cause minimal disruption but the 19-AA and especially 17-AA mutagenesis reveals a clear pattern. This is also observed for constructs O75056, Q13591, Q6UXE8, Q8IYV9, Q9H3T3, Q9NVM1, Q9ULG6 in the profiles of all proteins, provided in supplementary Fig. S4. In some cases, however, the mutational profiles are essentially identical across the three lengths. Western blot analysis of selected constructs at different lengths does not indicate that the phenomenon is related to their expression level (supplementary Fig. S6). Interestingly, this is not necessarily dependent on the reconstructed GFP fluorescence of the WT constructs, which can be very similar for the three lengths or differ wildly (e.g., Q8N6P7 vs Q8NFY4 and Q6UW88, supplementary Fig. S4). Remarkably, however, if the significant disruption patterns are observed in more than one TM length, these patterns are always consistent, as evident in Fig. 4a and also in all constructs in Fig. 5. This suggests that while different length constructs may behave differently with respect to their tendency to display disruption, the underlying interactions between the helices tend to be consistent.

**Fig. 5.**
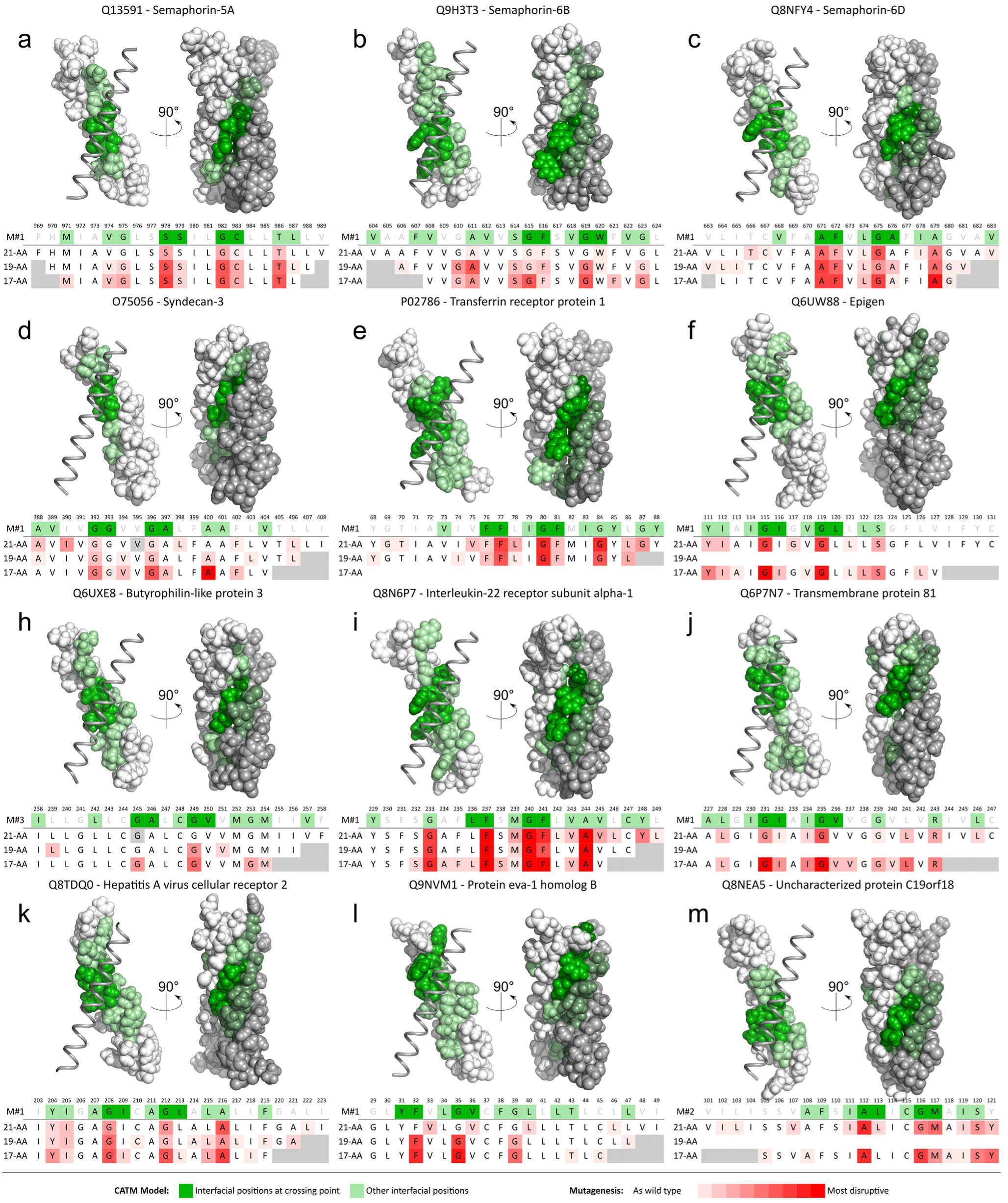
The mutagenesis of twelve constructs is consistent with the predicted GAS_right_ interface. The figure illustrates the structural model, with the interfacial positions highlighted in green. The darker green identifies the four residues that mediate the tight interactions at the crossing point between the helices. The same color-coding is provided in the sequence. M#1, M#2 and M#3 refer to CATM’s first, second and third top model, chosen for their matching with the mutational profile. The average profile of the three length constructs (21-AA, 19-AA and 17-AA) is show with the degree of disruption indicated in red.

### Mutagenesis supports the model of twelve predicted GAS_right_ homodimers

The twelve GAS_right_ dimers validated by the mutagenesis are illustrated in Fig. 5. They include proteins involved in important biological functions in processes, such as cell-surface signaling, proliferation and adhesion (semaphorins 5A, 6B and 6D, syndecan-3), transport (transferrin receptor protein 1) and the modulation of immune response (interleukin-22 receptor subunit alpha-1, butyrophilin-like protein 3, hepatitis A virus cellular receptor 2). The computationally predicted interfacial positions are highlighted in green in the structural representation and the sequence. A darker shade of green identifies the most central four amino acids that are involved in the tightest interaction at the point of closest approach between the helical pairs. These positions include GxxxG or GxxxG-like motif, with the Gly residues expected to be the most sensitive positions to mutation. Some of these proteins were previously known to form homodimers (all semaphorins, syndecan-3 and the transferrin receptor protein 1), but of these, the TM domain is known to mediate dimerization only for syndecan-3. There is no prior evidence of self-association for the other seven proteins (although some are known to form heterologous complexes with other proteins), therefore these findings may be helpful to shed light on their biological mechanisms.

To visualize the TM domain in the context of the entire protein, we also produced predictions using AF3. The TM domains of these predictions are compared to the CATM models in Fig. 6. In nine of the twelve cases, the models generated by the two programs are in excellent agreement with each other. In two cases, AF3 places the TM domains in proximity but not in contact or poorly packed. Interestingly, in one case (Butyrophilin-like protein 3), AF3 predicts with high confidence a well-packed helical dimer but its structure is different than the CATM model.

**Fig. 6.**
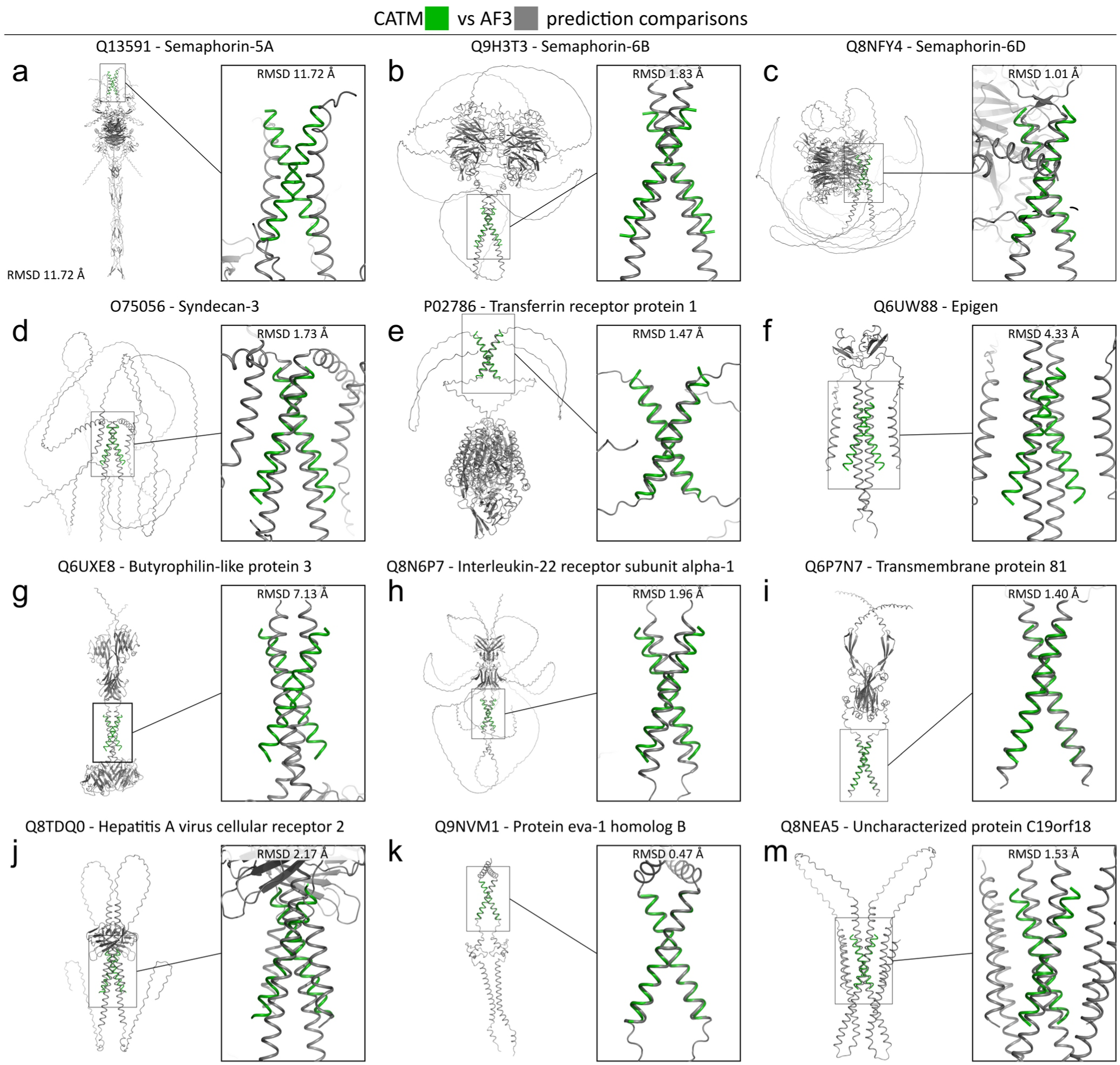
Correspondence between the CATM and AlphaFold3 models for twelve validated GAS_right_ constructs. The prediction from CATM is represented in green. The AF3 predictions are represented in gray.

A remarkable finding of the AF3/CATM comparison is that the level of correspondence between CATM and AF3 is very high only among this set of validated GAS_right_ constructs (9 out of 12, Table S3) whereas AF3 and CATM agree only in 3 of the remaining 88 constructs (3.4%). This disproportion is highly significant statistically (χ^2^ test, *p* = 4.4×10^-10^). The converging outcomes between an AI-based method that uses co-evolutionary information and a physics-based method increase the confidence that these structural predictions correspond to biologically relevant structures.

### Semaphorins 5A, 6B and 6D

The 100 TM helix library contained three members of the semaphorin (SEMA) family (SEMA 5A, 6B and 6D) and, remarkably, all three displayed strong TOXGREEN signals and mutagenesis profiles consistent with the predicted GAS_right_ model (Fig. 5a, b and c). Semaphorins are a diverse family of cell-surface signaling proteins originally discovered for their role in guiding axon development^68^ but are critical for regulating cell morphology, differentiation and cell-cell interactions in both neuronal and non-neuronal cell types, including in angiogenesis and immune response. Semaphorins engage in short-range cell-to-cell interactions with plexin receptors in adjacent cells^69^. These *“trans”* (between cells) interactions are mediated by semaphorins’ characteristic *sema* domain, which forms homodimers that bring together two plexin molecules, triggering their signaling^70–73^. The response to the signals typically induces significant cellular changes in the cytoskeletal and adhesive elements that specify cell morphology^74–76^, affecting integrin function and actin dynamics^77,78^, inducing actin filament disassembly^79^, and modifying membrane dynamics such as endo- and exocytosis^80^. Semphorins/plexin interactions can also occur in *“cis”*, between molecules on the same cells, and these are important to modulate or suppress intercellular signaling^81^.

SEMA 5A regulates neural circuit development in vertebrates^71,82,83^. In humans, it has been identified as an autism susceptibility gene^84,85^, and, in mice, SEMA 5A mutants negatively display increased numbers of excitatory synapses and deficits in social interaction, a hallmark of autism-spectrum disorders^86^. Additionally, SEMA 5A promotes angiogenesis in endothelial cells^87^. Structurally, SEMA 5A forms disulfide-linked dimers through its extracellular thrombospondin type-1 repeats (TSR)^88^, a unique feature of the semaphorin class 5 family that enables their interactions with the glycosaminoglycan moieties proteoglycans^89^. The crystal structure of SEMA 5A shows that its third and fourth TSR repeats link the two chains through a domain-swapped fold^90^. Our mutagenesis confirms that the TM helix SEMA 5A forms stable GAS_right_ dimers through an interface that involves residues M_971_xxVGxxSSxxGCxxT_986_ (Fig. 5a). The data therefore suggests that two SEMA 5A chains form an elongated dimeric structure interacting at the N-terminus through the sema domain, through the TSR domains in the middle of the protein, and via a GAS_right_ motif in the C-terminal TM domain.

The other two semaphorins (SEMA 6B and 6D) are members of the type 6 family, which is involved in neuronal and non-neuronal morphogenesis, primarily through interactions with class A plexins^91,92^. SEMA 6B is known as the receptor of a toxin released by the clostridium Paeniclostridium sordellii, which causes an almost invariably lethal toxic shock syndrome associated with gynecological infections^93,94^. SEMA 6D is involved in cardiac morphogenesis^95^, bone homeostasis, and immune response^96^. SEMA 6D is an interesting potential drug target since it can hamper cytotoxic T cell infiltration in head and neck cancer, therefore its modulation could increase the efficacy of immune checkpoint inhibitors for tumors resistant to immunotherapy^97^.

A characteristic of the semaphorin 6 family, is their long cytoplasmic C-terminal region^98^, which is critical for their ability to perform reverse signaling, i.e., transmitting external signals to their own cell through the recruitment and activation of kinases^99^. In SEMA 6B, mutations leading to truncation of this cytoplasmic region cause progressive myoclonic epilepsy in humans and produce defective development of brain neurons in zebrafish^100^. In SEMA 6D, reverse signaling regulates anti-inflammatory polarization and lipid metabolism in macrophages^101^ and is also important in chickens for cardiac development^99^.

The mutagenesis profile of both SEMA 6B-TM and 6D-TM are in good agreement with the CATM model. The profile of SEMA 6B identifies the positions most sensitive to mutation as the A_611_xxxGxxxG_619_ motif at the center of the predicted interface (Fig. 5b). The profile of SEMA 6D is also in excellent agreement with a predicted interface where A_671_FxLGAxxA_679_ are the most important residues (Fig. 5c).

The CATM models of SEMA 6B and 6D are confirmed by AF3 (1.73 and 1.01 Å RMSD, respectively), as illustrated in Fig. 6b and c. Interestingly, SEMA 5A is one of the two cases among the twelve validated GAS_right_ dimers for which AF3 does not predict an interaction between the TM helices (Fig. 6a). Overall, the computational and mutational data provide strong evidence that the TM helix of these semaphorins is a dimerization domain. Considering the frequent involvement of TM helices in mediating receptor signals in and out of the cell^3–5^, the sensitive positions identified by the validated interfaces provide the opportunity for assessing whether the TM dimers act as functional elements in these semaphorins.

### Syndecan-3

Syndecans are a family of four proteins that are responsible for cell signaling, proliferation, and adhesion in mammals. Their extracellular N-terminal region is covalently linked to glycosaminoglycan chains, which mediate interactions with a multitude of ligands, including growth factors, chemokines, cytokines, proteinases, adhesion receptors, and collagens^102,103^. The interactions between the cytoplasmic domain of syndecans and intracellular kinases and the actin cytoskeleton regulate cell adhesion and migration, with implications in both tumor progression and suppression^104^. All four syndecans homodimerize and form heterologous complexes among each other through interactions mediated by the GxxxG motif in their transmembrane domain^105,106^. Our mutational analysis included syndecan-3, which is expressed in neural tissues and developing musculoskeletal tissues. The mutagenesis confirms that syndecan-3 associates through a GAS_right_ dimer centered around its G_392_xxxGxxxA_400_ motif (Fig. 5d). The AF3 model of syndecan-3 is overall very low confidence but the program identifies a GAS_right_ motif similar to that predicted by CATM (1.73 Å RMSD, Fig. 6d).

### Transferrin receptor protein 1

The Transferrin receptor protein 1 (TfR1, Fig. 5e) mediates the cellular uptake of iron. Insoluble iron is distributed through the organism as Fe^3+^ bound to transferrin. Endocytosis of the TfR1/transferrin complex leads to the release of iron in the endosomal compartment through acidification, followed by the recycling of the apotransferrin-receptor complex to the cell surface ^107^. The receptor is necessary for the development of erythrocytes, and the immune and nervous systems and patients with TfR1 mutations can have immunodeficiency characterized by impaired T and B cell function^108^. The extracellular domain of TfR1 is a homodimer in solution^109,110^ therefore our data indicate that the TM domain is an additional dimerization domain for TfR1. The CATM model of the TfR1 TM dimer is in good agreement with the AF3 dimeric prediction (1.47 Å RMSD), which also models the transferrin binding domain connected to the TM domain by an extended low confidence region (supplementary Fig. 6e).

### Butyrophilin-like protein 3 (BTNL3)

Butyrophilin-like protein 3 (BTNL3, Fig. 5g) is one of five members of the human butyrophilin-like protein family. They are poorly understood proteins with co-stimulatory and inhibitory roles in the immune response^111,112^. BTNL3-TM displayed limited disruption in its mutagenesis profile, with a pattern emerging only in the 17-AA construct, highlighting the sensitivity of the G_245_xxxGxxxG_253_ motif. Interestingly, AF3 produces a model in which the interaction interface is diametrically opposite to this GxxxGxxxG motif (Fig. 6g). The AF3 model is high-confidence and well packed, however, its interfacial positions (L_240_xxxCxxLxxxVxxM_254_) are not susceptible to mutation in our TOXGREEN analysis. The presence of two potential interaction interfaces, however, may explain the limited sensitivity to mutation. If these two configurations are physiological, it suggests that BTNL3’s function somehow involves a conformational switch in its TM region, corresponding to a nearly 180° rotation around its TM helices.

### Interleukin-22 receptor subunit alpha-1 (IL-22R1)

IL-22R1 and IL-22R2 form a heterodimeric transmembrane receptor complex that binds to the interleukin 22 cytokine and activates a number of immune cell types^113^. There are currently no reports in the literature indicating that IL-22R1 can form homodimeric complexes. The mutagenesis of IL-22R1-TM (Fig. 5h) displays similar profiles across the 21-, 19- and 17-AA constructs, with strong disruption at positions G_233_xxxFxxGFxxA_244_, in excellent agreement with CATM’s molecular model. The AF3 prediction is also in agreement with CATM (RMDS 1.96 Å, Fig. 6x).

### Transmembrane protein 81 (TMEM81)

TMEM81 was recently discovered to be part of a trimeric complex formed with two other single-pass fertilization factors, Izumo1 and Spaca6, which are essential in the binding and fusion of sperm and egg^114^. In this recent study, the authors produced an AlphaFold multimer model of the Izumo1/Spaca6/TMEM81 complex. In that model, the TM domains of Izumo1 and TMEM81 form a GAS_right_ heterodimer using the same positions (A_227_LxxGIxxGVxxG_239_) that are involved in the homodimer predicted by CATM (Fig. 5i). The AF3 homodimeric model of TMEM81 is in excellent agreement with CATM’s model (RMDS 1.40 Å, Fig. 6i). If both the homodimeric and heterodimeric forms of TMEM81 are physiological, they would have to be alternative states occurring at different functional or assembly stages of these proteins.

### Hepatitis A virus cellular receptor 2 (HAVCR2)

HAVCR2, also known as TIM-3, is a cell-surface receptor implicated in modulating innate and adaptive immune responses in T-cells and other immune cells. TIM-3 is recruited to the immunological synapse^115,116^. It is known to bind to several ligands, which induce the phosphorylation of tyrosine residues in its cytoplasmic tail^117^. This triggers the dissociation of BAT3 adaptor protein and association with the FYN kinase, leading to the inhibitory activity on T cell receptor signaling^118^. Our mutagenesis indicates that TIM-3-TM homodimerizes through an interface formed by Y_204_IxxGIxxxGLxxLA_216_ (Fig. 5j). The prediction is confirmed by AF3, which is relatively similar and utilizes the same interface (RMDS 2.17 Å, Fig. 6j).

### Other proteins

**Protein eva-1 homolog B** (EVA1B) is a poorly studied member of the EVA1 family. Its better known paralog EVA1A localized in the endoplasmic reticulum and lysosome and it is involved in regulating autophagy and apoptosis^119^. EVA1B is upregulated in glioma and is considered a poor prognostic biomarker in this disease^120^. We found that EVA1B-TM oligomerizes strongly in TOXGREEN in a manner that is highly sensitive to mutation in its F_32_xxGxxxG_39_ motif (Fig. 5k). The AF3 model is nearly identical to the CATM model (RMDS 0.47 Å, Fig. 6k). **Epigen** is a small single-pass membrane protein. Following proteolysis, its extracellular domain acts as an Epidermal growth factor receptor ligand, promoting cell differentiation^121,122^. We found no reports indicating epigen homodimerization. CATM’s prediction of Epigen-TM (Fig. 5f) is not replicated by AF3, which produces two poorly interacting parallel helices (Fig. 6f). **C19orf18** is a protein of unknown function (Fig. 5m) that is up-regulated in pancreatic cancer^123^. CATM and AF3 predict similar models (RMDS 1.53 Å, Fig. 6m).

Finally, three of the mutagenesis profiles identified here do not match the CATM predictions but are consistent with their AF3 models (illustrated in supplementary Fig. S5). These are **Oxidized low-density lipoprotein receptor 1**, which is implicated in organ damage caused by dyslipidemia and that mediates the internalization and degradation of oxidatively modified low density lipoprotein by endothelial cells^124^. The second is **Cytotoxic T-lymphocyte protein 4**, a receptor that acts as a negative regulator of T-cell responses^125^. Lastly, **Cell cycle progression protein 1** is ER-resident protein that is transcriptionally upregulated upon ER stress and is important for stimulating ER selective autophagy^126^.

## Conclusions

TOXCAT is a widely employed assay for the determination of the oligomeric properties of TM helices. It has contributed to our understanding of the biology and the folding of numerous single-pass membrane protein systems. Here we demonstrate that its GFP-expressing variant, TOXGREEN, is applicable to the simultaneous analysis of oligomerization in large libraries of TM helices. This rapid, large-scale evaluation opens the door to new experiments for understanding the interactions, energetics and biology of single-pass protein systems as well as new protein engineering approaches.

We have shown that the reconstructed fluorescence values obtained statistically from FACS and NGS data reproduce the direct measurements of individual constructs with good accuracy and sensitivity. We also established a maltose growth test compatible with large libraries, which is necessary for identifying constructs that may not be inserted correctly or efficiently in the membrane. One aspect of the workflow that cannot be performed yet on a large scale is the systematic assessment of protein expression, which is typically performed using western blots. To address this issue, Fleishman and coworkers previously replaced the MBP moiety of the TOXCAT construct with the antibiotic resistance enzyme β-lactamase and used selection in ampicillin as a proxy for expression and insertion^30^. Since the TOXGREEN and TOXCAT constructs are identical (the only difference being the nature of the reporter gene), the same strategy could be adopted in TOXGREEN sort-seq. It should be noted that controlling for expression is not necessary for obtaining quality mutagenesis profiles because closely related constructs, such as point mutants, tend to have similar levels of expression^27,42,127^.

TOXGREEN sort-seq produced high-quality, reproducible mutational profiles from a large library of 17,400 constructs. We found that the longer TM inserts, and particularly the 21-AA library, produced stronger fluorescence signals but were less susceptible to disruption. The shorter constructs tend to fluoresce more weakly, but they have a higher propensity to display disruption patterns when they produce sufficient signal. Based on these observations, we did not identify an optimal insert length and suggest that testing each TM helix at multiple lengths may be an effective way to address this issue. Our large-scale analysis also confirmed that the “limited” strategy of three mutations at each position can be effective in determining an interaction interface, although a larger selection of mutants (such as A, I, F, G, L, M and V) is likely to be more informative when affordable.

TOXGREEN sort-seq can be used to rapidly evaluate the dimerization propensity of entire groups of single-pass proteins, to study their energetics or to engineer TM dimers. The present analysis was used as an opportunity to further assess CATM’s performance in predicting GAS_right_ homodimers from primary sequence. The algorithm performed well when tested against the small set of known GAS _right_ dimers^43^, but its propensity for producing false positives was not previously assessed because of the limited availability of single-pass proteins in the structural database. In the present analysis, 35 constructs produced a weak TOXGREEN signal and 31 did not display an appreciable disruption pattern. Of the remaining 34 constructs, CATM’s GAS_right_ predictions were supported in 12 cases (35%). Remarkably, this subset has a high rate of correspondence with the AF3 predictions, supporting the hypothesis that the models correspond to biologically relevant structures. The high-throughout mutagenesis data provided by TOXGREEN sort-seq could be used in the future to improve the algorithm’s precision. It should also be noted that 13 constructs produced a pattern that mapped to a possible alternative interface. Three of these mutagenesis profiles are corroborated by the AF3 prediction, suggesting that these interfaces are also biologically relevant.

## Methods

### Computational prediction of dimeric GAS_right_ structures

The structural models of the 100 GAS_right_ dimers were predicted in previous work from all human protein sequences annotated as single-pass membrane proteins in Uniprot using CATM^42^. These models are available at https://catm.biochem.wisc.edu/CATM. As summarized in supplementary Table S4, the sequences were selected among models with a CATM energy score of at least -10 kcal/mol. For consistency, we limited the selection to sequences with a predicted length of 21 amino acids, which is the most common length annotated for transmembrane domains in Uniprot. Sequences containing proline were excluded because their tendency to create kinks in α-helices. Sequences containing more than one strongly polar amino acid (D, E, H, K, N, or Q) were also removed because these amino acids have a tendency to cause non-specific association in TM helices driven by hydrogen bonding^67^. Any sequences for which part of the predicted interface included any residues outside of the TM domain were also removed. The final library consisted of 100 biological TM helices expressed at three different lengths, a 21 amino acid full-length construct and truncations to 19 and 17 amino acids operated at the N- and/or C-terminal ends of the helices, eliminating positions predicted not to be involved at the dimerization interface. The final library thus consisted of 300 WT sequences, plus three mutants for each position at each length, for a theoretical total of 17,400 constructs (100 constructs ×(3 WT + 3×21 + 3×19 + 3×17 mutants)).

The full-length dimeric structures of each candidate GAS_right_ protein were also predicted using AlphaFold 3^128^ via the AlphaFold Server available at https://alphafoldserver.com/. The full sequence of every protein was used as input with two copies of every sequence for multimer prediction. All parameters were set as default.

### Cloning and Expression of Chimeric Protein Libraries

All sequences encoding for TM domains of interest were ordered as oligo pools from Twist Bioscience. Distinct predefined subsets (segments) of the oligo pools were individually amplified using unique primer pairs specific to each segment by quantitative PCR (KAPA SYBR Fast qPCR master mix, Roche) based on the protocol outlined in Kosuri et al.^129^ followed by another traditional PCR reaction using the qPCR reaction as a template. These amplified segments were digested with NheI and DpnII and cloned into the pccGFPKan vector^32^ at the NHeI-BamHI restriction sites. The DNA sequences of the inserts were designed such that the last two nucleotides are “TT” before the DpnII cut site “G” resulting in an in-frame “L” once ligated into the corresponding pccGFPKan restriction sites. The resulting chimeric protein sequences are: RAS–TM Sequence of Interest–LILI (e.g., RAS– LLAAGILGAGALIAGMCFIII–LILI for LLAAGILGAGALIAGMCFIII). TOXGREEN construct library segments were transformed into *malE* deficient *E. coli* strain MM39. Each transformation outgrowth was grown overnight at 37 °C in 3 mL LB containing 100 μg/mL ampicillin to store segments or libraries as 25% glycerol stocks. Segments were determined to be successfully cloned if the calculated colony-forming units on LB agar plates supplemented with 100 μg/mL ampicillin were at least tenfold the number of sequences in the segment. To create composite libraries from individual segments or colonies, each library component was inoculated into 3 mL LB containing 100 μg/mL ampicillin and grown overnight at 37 °C. 30 uL of each overnight culture was used to inoculate 3 mL fresh LB containing 100 μg/mL ampicillin and incubated at 37 °C until an optical density of approximately 0.1 at 600 nm was reached. The optical density at 600 nm of each culture was normalized by theoretical unique sequence count and then combined in equal proportion for every component to a final volume of 1 mL. 25% glycerol stocks were prepared for all combined libraries for storage at -80 °C.

### Selection of individual constructs for the validation set

A “validation set” of approximately 100 constructs was created for the purpose of developing the TOXGREEN sort-seq method (supplementary Table S1). The mutation length libraries were sorted using a SONY LE-SH800 into three populations: low, medium, and high fluorescence. The low bin was defined with any fluorescence lower than the median fluorescence of the monomeric mutant of GpA, G83I. The medium bin was drawn with the lower bound set to the upper bound of the low bin and the upper bound set to the median fluorescence value of GpA. The high gate was set to any cells with fluorescence above the median fluorescence level of GpA. The sorted cells were plated on LB supplemented with 100 μg/mL ampicillin and incubated at 37°C overnight. A total of 109 colonies were chosen between each of these pools and sent for sequencing. This resulted in 98 confirmed variants and the standard three traditional controls of GpA, GpA-G83I, and the base No-TM pccGFPKan plasmid. TOXGREEN fluorescence was measured individually for each construct in the set.

### MalE Complementation Assay

To confirm proper membrane insertion and orientation of individual TOXGREEN constructs, overnight cultures were plated on M9 minimal medium plates containing 0.4% maltose as the only carbon source and grown at 37 °C for 72 h^27^.

MalE complementation assay for the libraries was performed using liquid M9 maltose minimal media. Overnight cultures of each library were used to separately inoculate 500 mL M9 maltose media to a final concentration of 0.00125 optical density at 600 nm. Samples were taken following 30-36 h of growth at 37°C. DNA was extracted from M9 media samples along with the overnight LB cultures used for initial inoculation and sent for NGS.

The negative (non-inserting) subset of constructs from the validation set (highlighted in supplementary Table S1) was included in any liquid MalE complementation assay sample as a threshold growth level for any failing samples. For all protein sequences retrieved from the NGS, the fractional change in population percentage was calculated between the 30-36 h M9 time point and the LB overnight culture sample. Any sample with a fractional change equal to or worse than the best-performing negative sample was removed from further analysis.

The sequences for the negative control set as well as two positive controls have been deposited on the AddGene repository with the following accession numbers: 1E01: AddGene: 239013; 1E11: AddGene: 239083; 2E11: AddGene: 239084; 2F12: AddGene: 239085; 2H11: AddGene: 239086; 1F02: AddGene: 239087; 2H07: AddGene: 239088.

### Traditional TOXGREEN

Standard plate reader-based sfGFP expression was quantified in stationary phase as described prior^32^. Briefly, a freshly streaked colony was inoculated into 3 mL of LB broth containing 100 μg/mL ampicillin and grown overnight at 37 °C for all samples. To reduce background, 1.5 mL of cells were collected by centrifugation at 17,000 g and concentrated by re-suspension in 0.5 mL in PBS solution (137 mM NaCl, 2.7 mM KCl, 10 mM Na_2_HPO_4_, 2 mM KH_2_PO_4_, pH 7.4). 300 μL of each sample was transferred to a 96-well black-walled, clear bottom plate (Fisher Scientific) for fluorescence measurement. Fluorescence measurements were performed using an Infinite M1000 Pro plate reader (Tecan), using an excitation wavelength of 485 nm and recording emission from 500 to 600 nm. The relative sfGFP expression was calculated by normalizing the fluorescence emission at 512 nm to the optical density of the sample at 600 nm measured on an Agilent Technologies Cary 60 UV-Vis.

### Flow cytometry reading of individual TOXGREEN clones

Flow cytometry-based sfGFP expression was measured using a SONY LE-SH800 with a SONY 70 μM chip. The forward scatter (FSC) threshold was set at 0.05%. The sensor gains for FL1 (GFP) and backscatter (BSC) were set at 90% and 30%, respectively. The first gate was drawn to select for cells by eliminating cell debris on a side scatter area (SSC-A) vs. forward scatter area (FSC-A) plot. The second gate was drawn to eliminate doublet cells on a forward scatter height (FSC-H) vs. FSC-A plot. A freshly streaked colony was inoculated into 3 mL of LB broth containing 100 μg/mL ampicillin and grown overnight at 37 °C for all samples. To reduce background, overnight cultures were added to a PBS solution at a dilution level such that the event rate was below a maximum of 10,000 events per second. Median GFP fluorescence was calculated from gated cells for each variant. The FCS files were analyzed using FlowJo.

### Immunoblotting for individual constructs

Expression level of the length variants of several constructs was assessed by Western blot analysis (Fig. S6). 2 mL of overnight culture were pelleted, resuspended, and lysed through chemical lysis (SoluLyse, AMSBIO). For each sample, 45 μL of lysate was added to 15 μL 4X LDS loading buffer (Novex, Life Technologies) with β-mercaptoethanol and heated to 70 °C for 3 min. Cell lysate protein concentrations were determined by bicinchoninic acid assay (Pierce). 10 μg of each LDS protein sample was loaded onto a NuPAGE 4-12% Bis-Tris SDS-PAGE gel (Invitrogen) to separate by SDS-PAGE and transferred to polyvinylidene difluoride membrane (VWR). Peroxidase-conjugated monoclonal anti-maltose binding protein antibodies (Sigma-Alridch) were used for immunoblotting analysis. Blots were developed with ECL Western Blotting Substrate Kit (Pierce) and imaged using an iBright FL1500 (Invitrogen).

### Fluorescence-activated cell sorting

50 μL of any library glycerol stock was used to inoculate 3 mL of LB broth supplemented with 100 μg/mL ampicillin and grown overnight at 37 °C. FACS was performed using a SONY LE-SH800 cell sorter with a 70-μm microfluidic sorting chip. The FSC threshold was set at 0.05%. The sensor gains for FL1 (GFP) and BSC were set at 90% and 30%, respectively. The first gate was drawn to select for cells by eliminating cell debris on a SSC-A vs. FSC-A plot. The second gate was drawn to eliminate doublet cells on a FSC-H vs. FSC-A plot. Every library was appropriately diluted in PBS buffer with a flow rate set such that the event rate was at most 10,000 events per second. A suitable number of gates (4-7) were drawn to span the fluorescence distribution of the entire library ensuring every bin was distinguishable from its adjacent bins when sorted. 500,000 events were sorted into every bin. All bins were sorted into 2 mL LB broth containing 100 μg/mL ampicillin. Sorted populations were grown in a total of 5 mL LB broth containing 100 μg/mL ampicillin until an optical density at 600 nm of approximately 0.1 and plasmid DNA was extracted.

### Library preparation and deep sequencing

For each sample for NGS, the sequences corresponding to the TM helices were amplified through PCR from the extracted TOXGREEN plasmid DNA followed up by another PCR reaction in order to add the necessary Illumina TruSeq adapter, unique barcode, and “stem” sequences. The products were submitted to the University of Wisconsin Biotechnology Center DNA Sequencing Facility for Index PCR (TruSeq) and Illumina NovaSeq6000 (2×150 shared) Sequencing services.

### Sequence Analysis and TOXGREEN fluorescence reconstruction

A minimum cutoff of 10 copies of a particular sequence had to be present in the NGS for it to be retained. After filtering, each DNA sequence was translated to amino acids and matched to its appropriate TM in length, protein ID, and mutation. All of the above analysis was performed with *ad hoc* Perl scripts. Statistical reconstruction calculations of sfGFP fluorescence output were performed in R and calculated as a weighted average. The distribution of each individual library construct in the bins is then used to reconstruct their estimated GFP fluorescence value with a statistical inference approach based on a weighted average, adapted from Kosuri et al.^57^. Briefly, for each construct *i* within each sorted bin *j,* we first calculate the normalized fractional contribution *a_ij_* as follows:

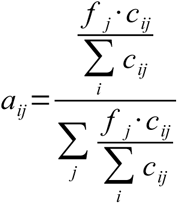

where *c_ij_* is the sequencing count from NGS for construct *i* in bin *j*, and *f_j_* is the fraction of the sorted sample population in bin *j*.

We then obtain the reconstructed fluorescence, expressed as Relative Fluorescence Units (*RFU_i_*) for each construct *i* by summing the product of the normalized fractional contribution *a_ij_* and the median fluorescence *m_j_* over each sorted bin *j*.

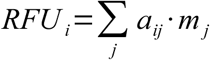

### Mutagenesis Analysis

The disruption level for every mutant was set based on its relative disruption compared to its WT reconstructed fluorescence. A minimum threshold of 50,000 RFU was set for performing the mutagenesis analysis to guarantee sufficient dynamic range. This value was established empirically and corresponds to approximately the 50% percentile in the distribution of RFU of all WT and mutant constructs (supplementary Fig. S7). For each mutant, a disruption value from 1 to 5 was calculated, with 1 being the most disruptive and 5 being similar or better than the WT. First, a quintile *RFU_Q_* was calculated as one-fifth of the difference between the reconstructed fluorescence of the WT *RFU_WT_* minus a base fluorescence value in the monomeric range *RFU_B_*.

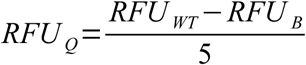

*RFU_B_* was chosen as 35,000, corresponding to approximately the 20% percentile in the distribution of all constructs (supplementary Fig. S7). Then the disruption index *DI_m,i_* of a mutant *m* at position *i* was calculated from its reconstructed fluorescence *RFU_m,i_* as the following:

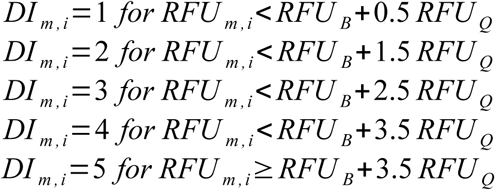

Finally, for each position *i*, an overall position-based disruption index *DI_i_* was calculated by averaging the disruption value of all *M* mutants (with typically *M* = 3) at that particular position:

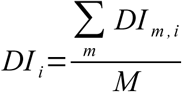

## Acknowledgments

This work was supported by National Institutes of Health grant R35 GM130339 to A.S. and National Science Foundation grant CHE-1710182 to A.S.. S.M.A. acknowledges the support of National Library of Medicine training grant T15 LM007359 to the CIBM Training Program and the Dr. James Chieh-Hsia Mai Wisconsin Distinguished Graduate Fellowship in Biochemistry. J.C. acknowledges the support of a Denis R.A. and Martha Washburn Wharton Fellowship in Biochemistry. M.L. acknowledges the support of National Institutes of Health training grant T32 GM08293 to the Molecular Biophysics Training Program. We are grateful to Dr. Kyle Nishikawa for helpful suggestions and discussion.

## Abbreviations

AF3: AlphaFold 3
BSC: backscatter
BTNL3: Butyrophilin-like protein 3
CAT: Chloramphenicol AcetylTransferase
EVA1B: Protein eva-1 homolog B
FACS: Fluorescence-Activated Cell Sorting
FSC: forward scatter
FSC-A: forward scatter area
FSC-H: forward scatter height
GFP: Green Fluorescent Protein
GpA: glycophorin A
HAVCR2: Hepatitis A virus cellular receptor 2
IL-22R1: Interleukin-22 receptor subunit alpha-1
MBP: Maltose Binding Protein
NGS: Next-Generation Sequencing
RFU: Relative Fluorescence Units
SEMA: semaphorin
SSC-A: side scatter area
TfR1: Transferrin receptor protein 1
TM: transmembrane
TMEM81: Transmembrane protein 81
TSR: thrombospondin type-1 repeats
WT: wild-type

## SUPPLEMENTARY INFORMATION

**Table S1.**
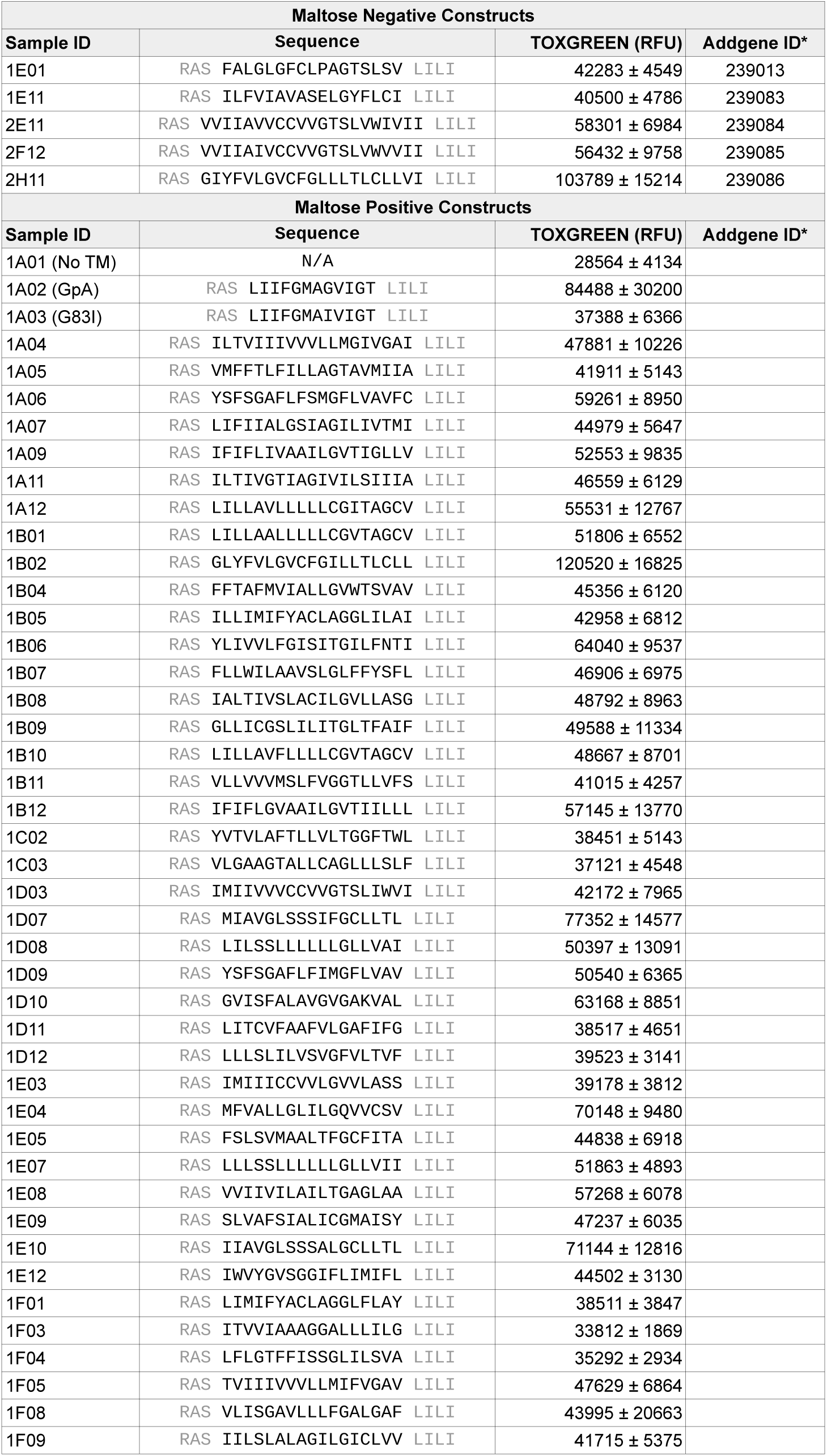
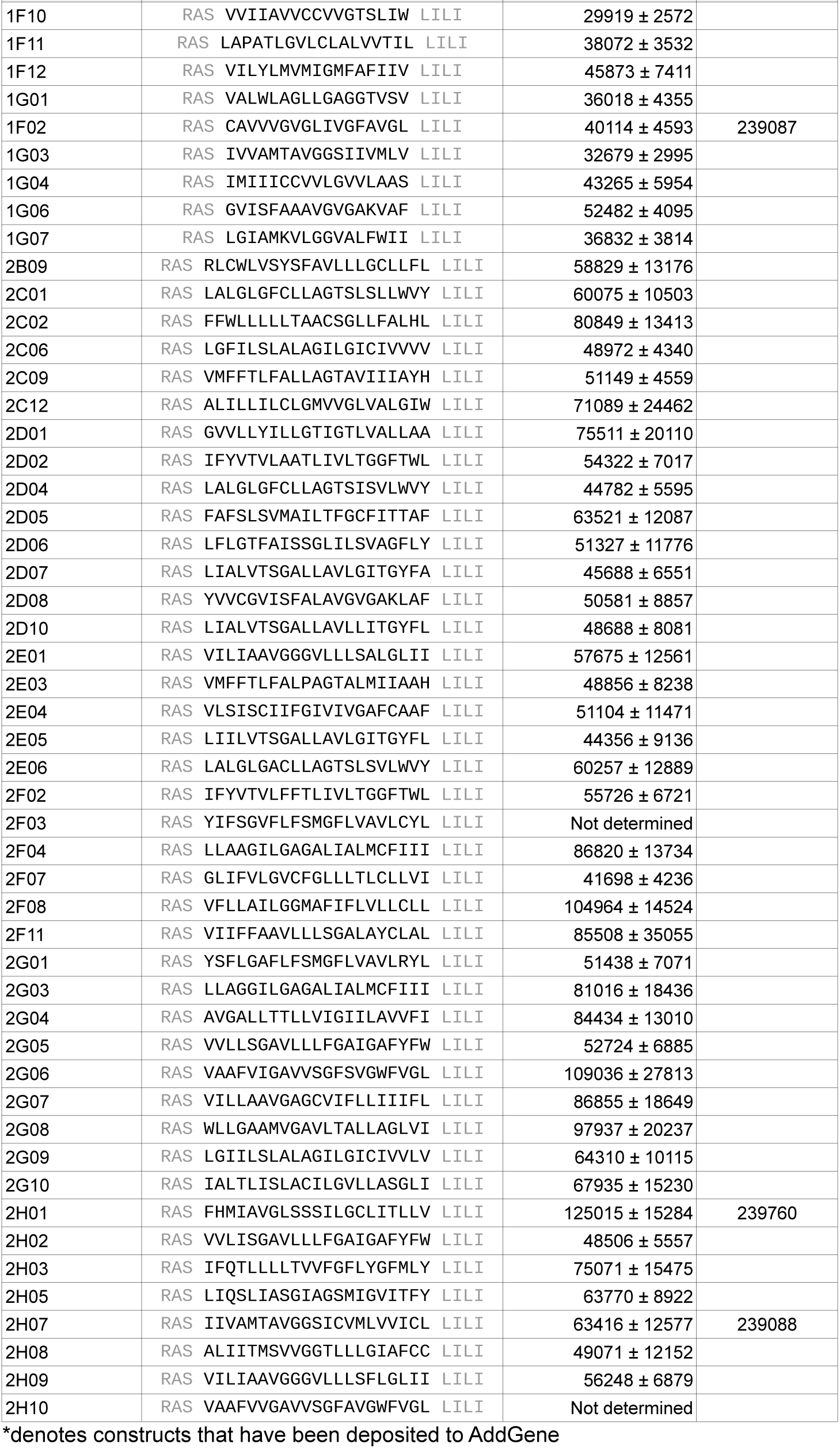
Set of 100 constructs used for the validation of the TOXGREEN sort-seq method (validation set).

**Fig. S1.**
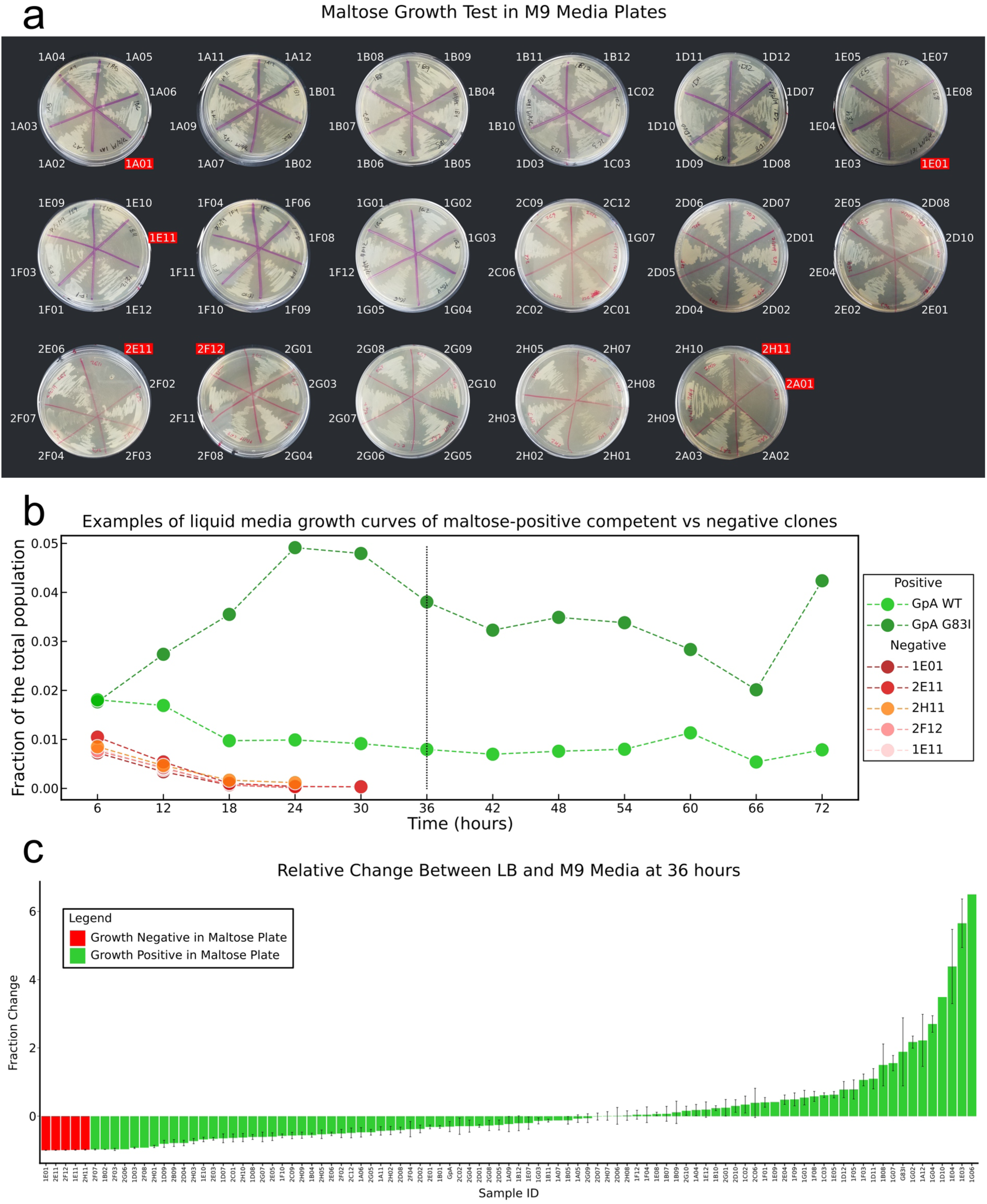
Adaptation of the maltose growth test for libraries in liquid minimal media. a) Traditional maltose growth test performed in M9 minimal media with maltose as the sole carbon source on each individual constructs of the validation library. The maltose-negative constructs are highlighted in red. b) Liquid media competition maltose growth assay. The figure illustrates the relative abundance of two maltose-positive constructs (shades of green) five maltose-negative constructs (shades of red). By 36 hours the maltose-negative constructs have disappeared from the colture. c) Fraction change between LB and M9 minimal maltose media for all constructs of the validation library. Error bars represent standard deviation of three replicates.

**Table S2.**
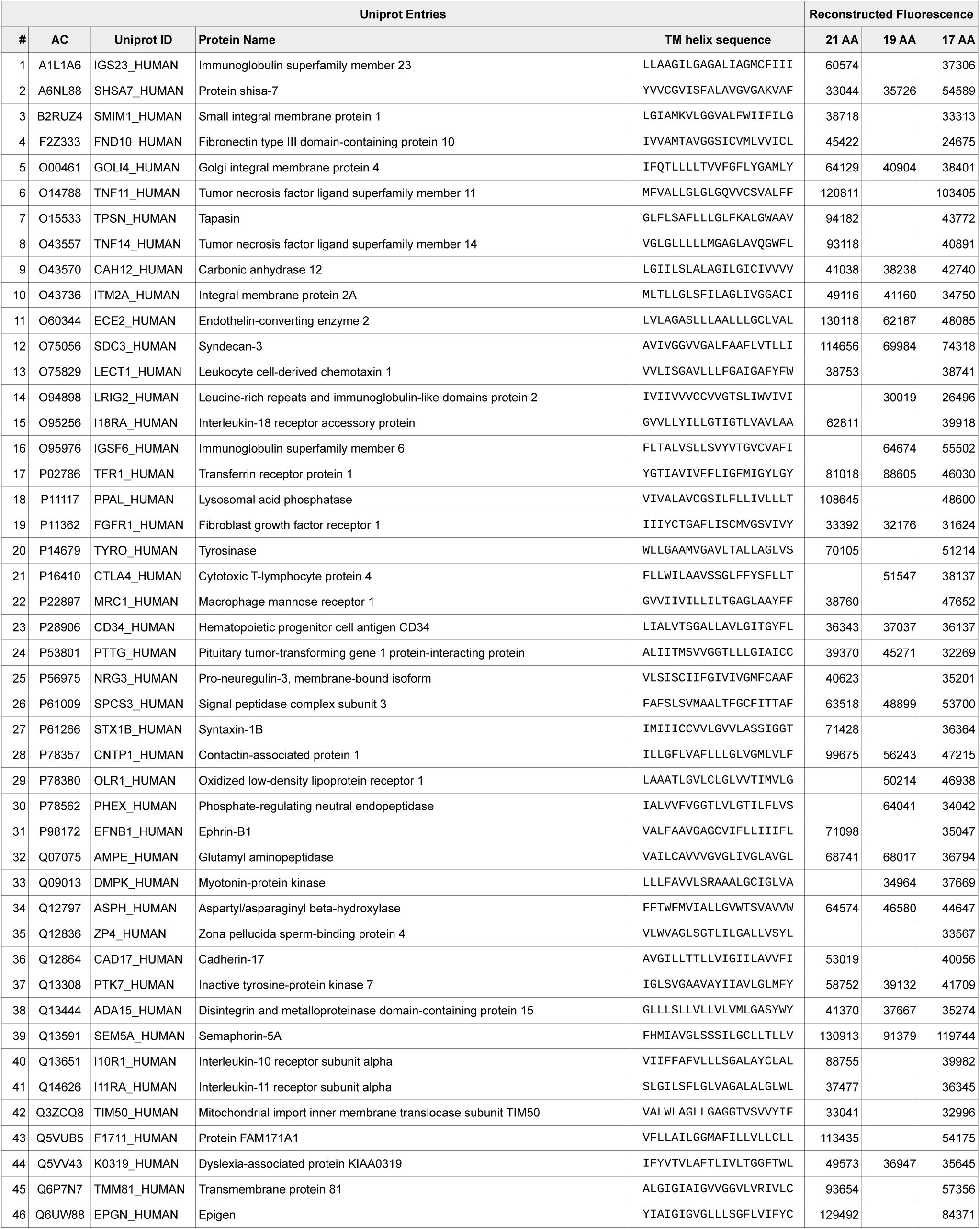
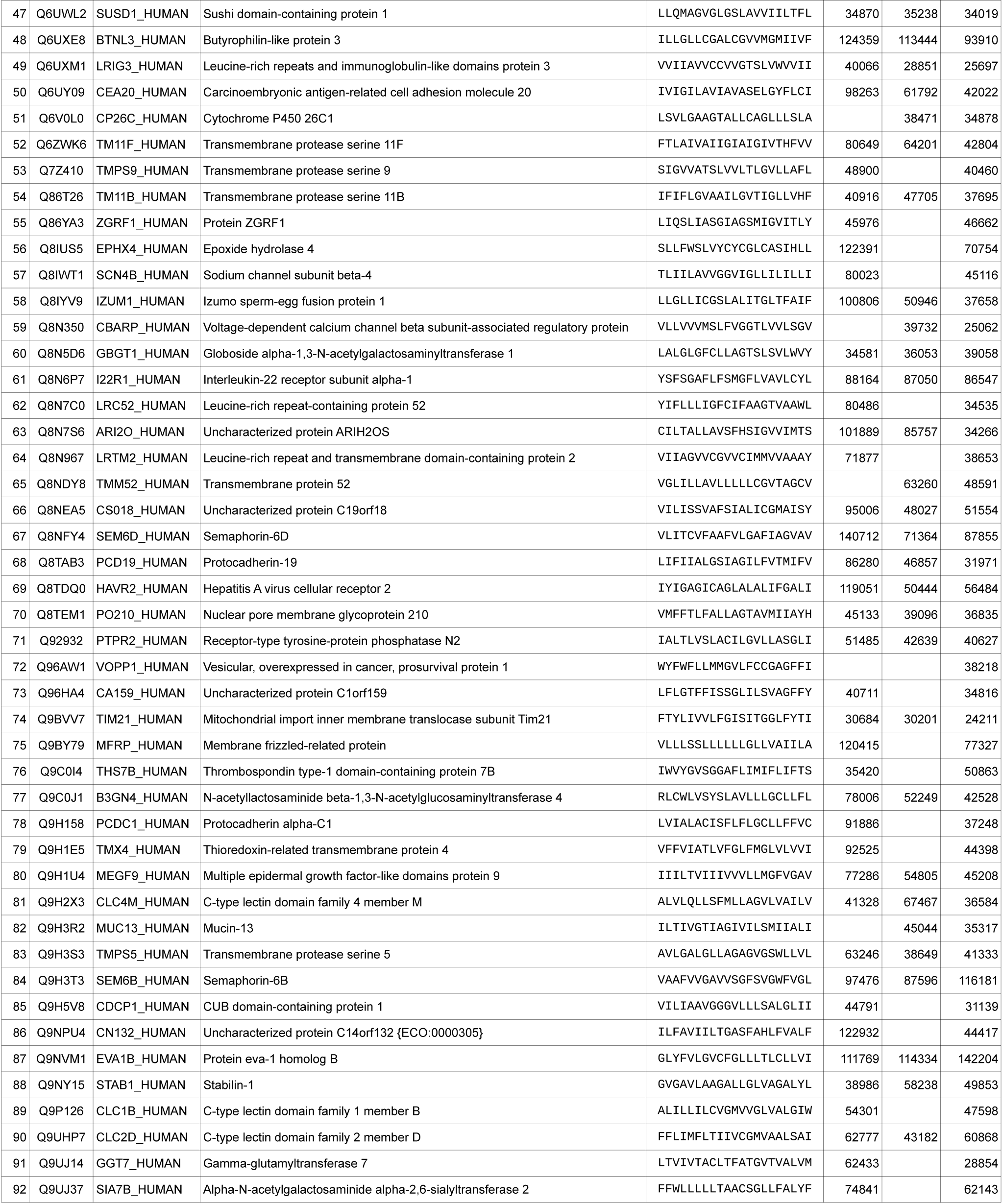
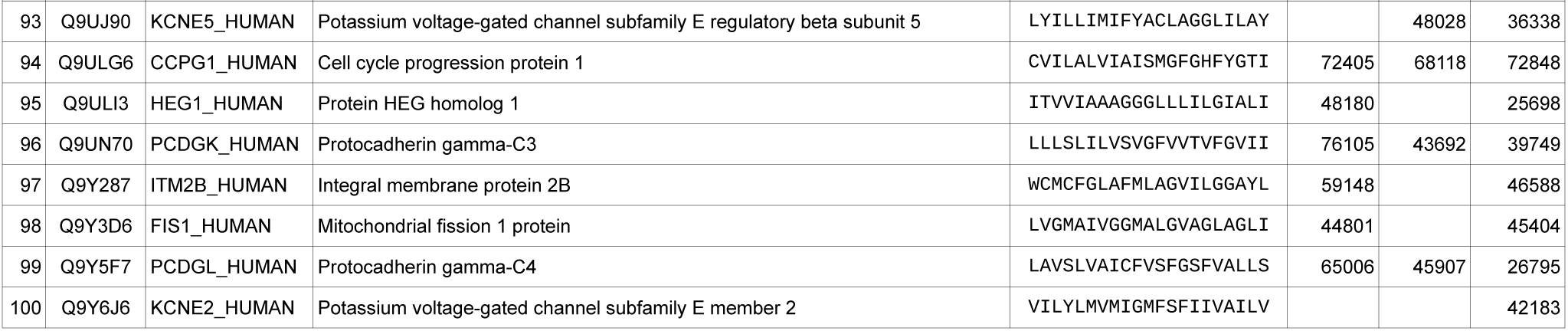
Library of 100 computationally predicted GAS_right_ dimers.

**Fig. S2.**
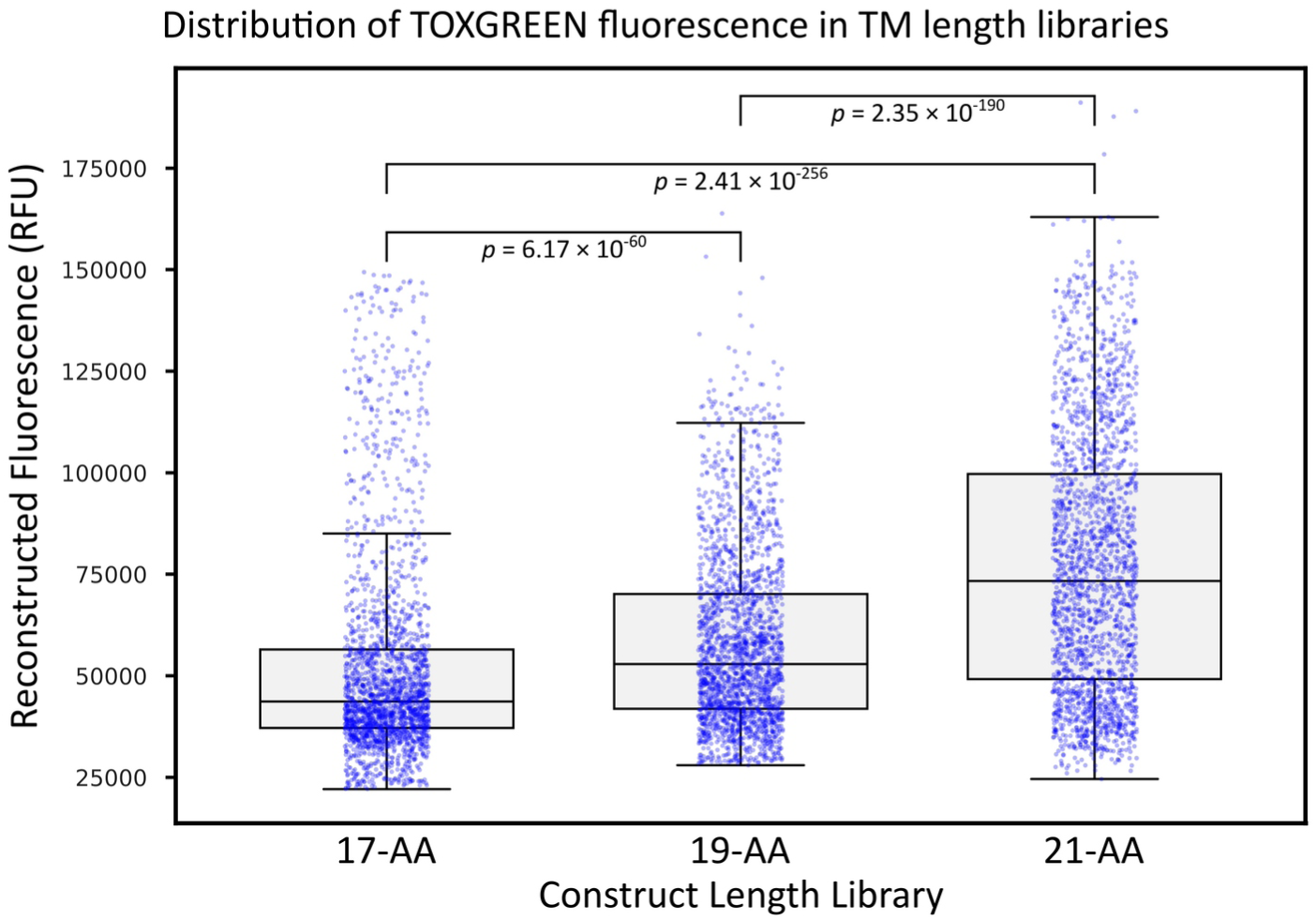
Distribution of reconstructed fluorescent signal for all constructs from libraries with different TM helix length. Statistical significance calculated with the Wilcoxon Signed-Rank Test.

**Fig. S3.**
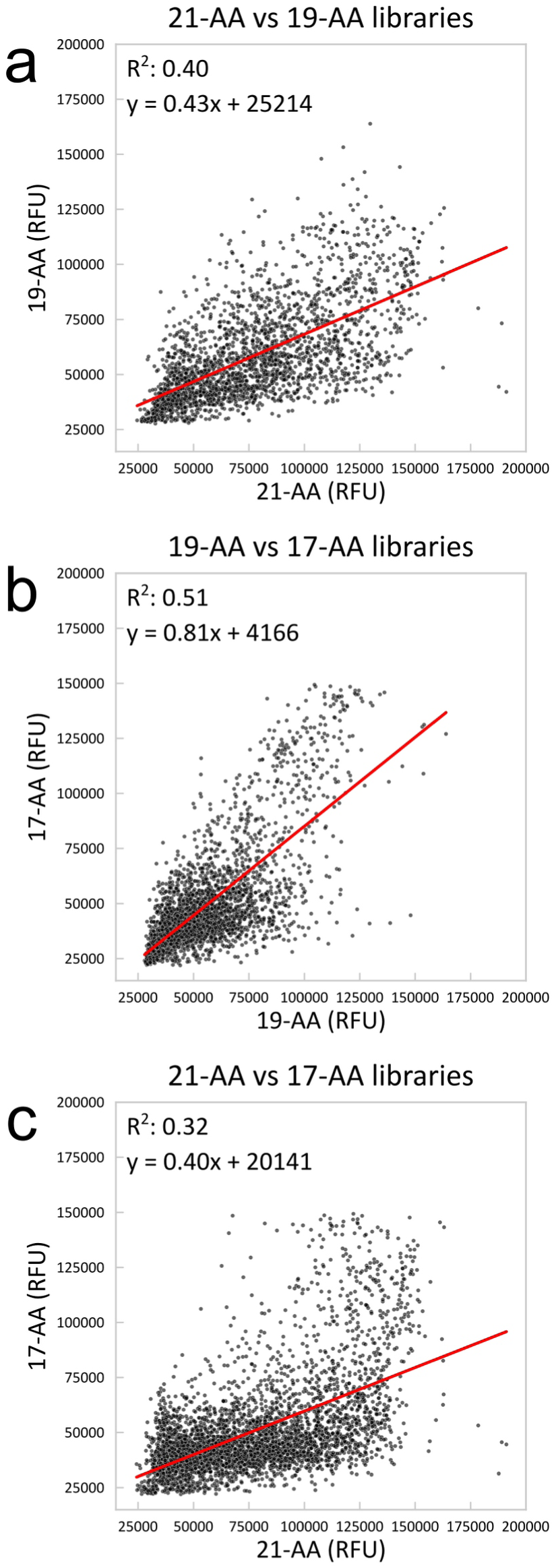
Correlation of the reconstructed fluorescence in the three length libraries. a) 21-AA vs 19-AA libraries. b) 19-AA vs 17-AA libraries. c) 21-AA vs 17-AA libraries.

**Fig. S4.**
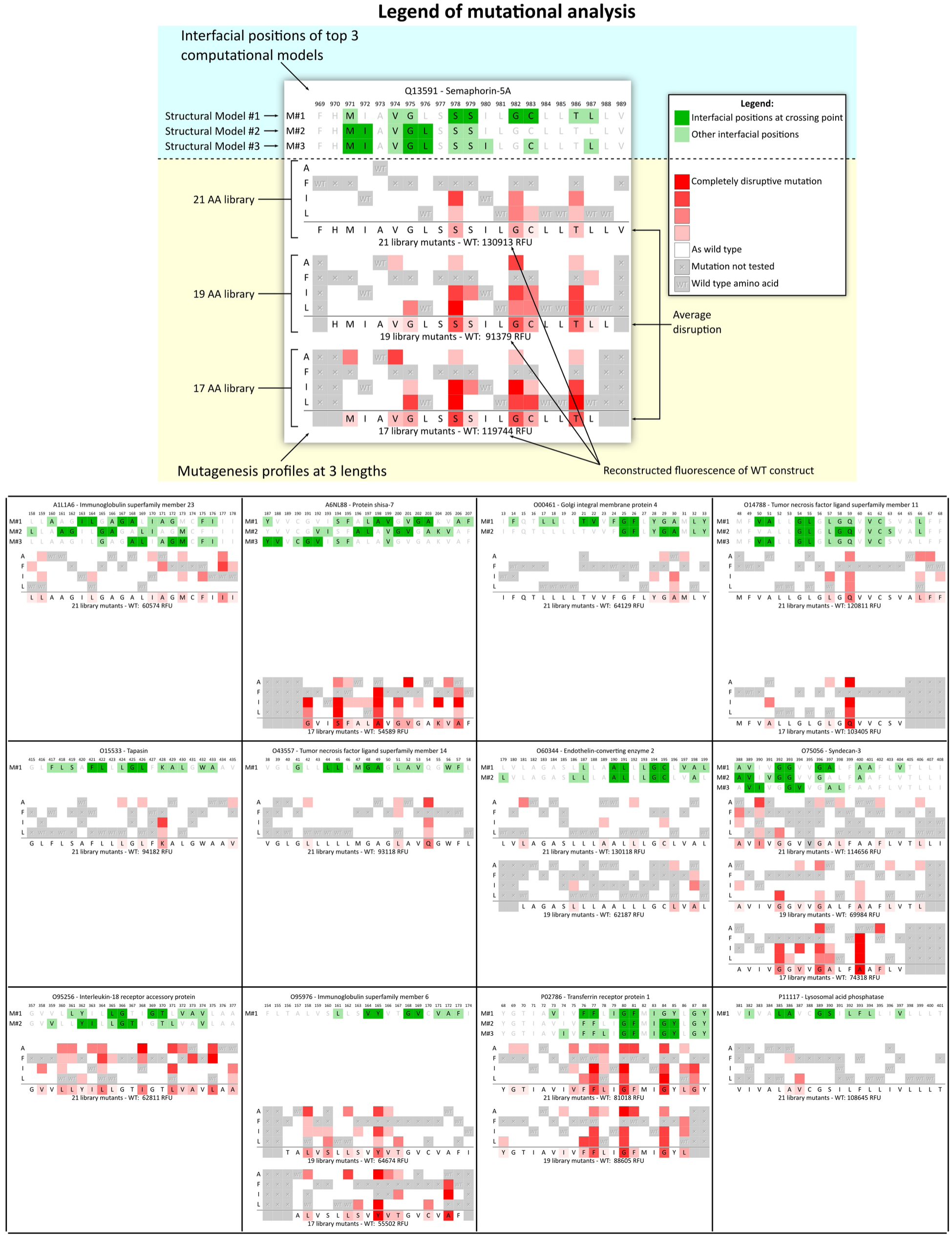
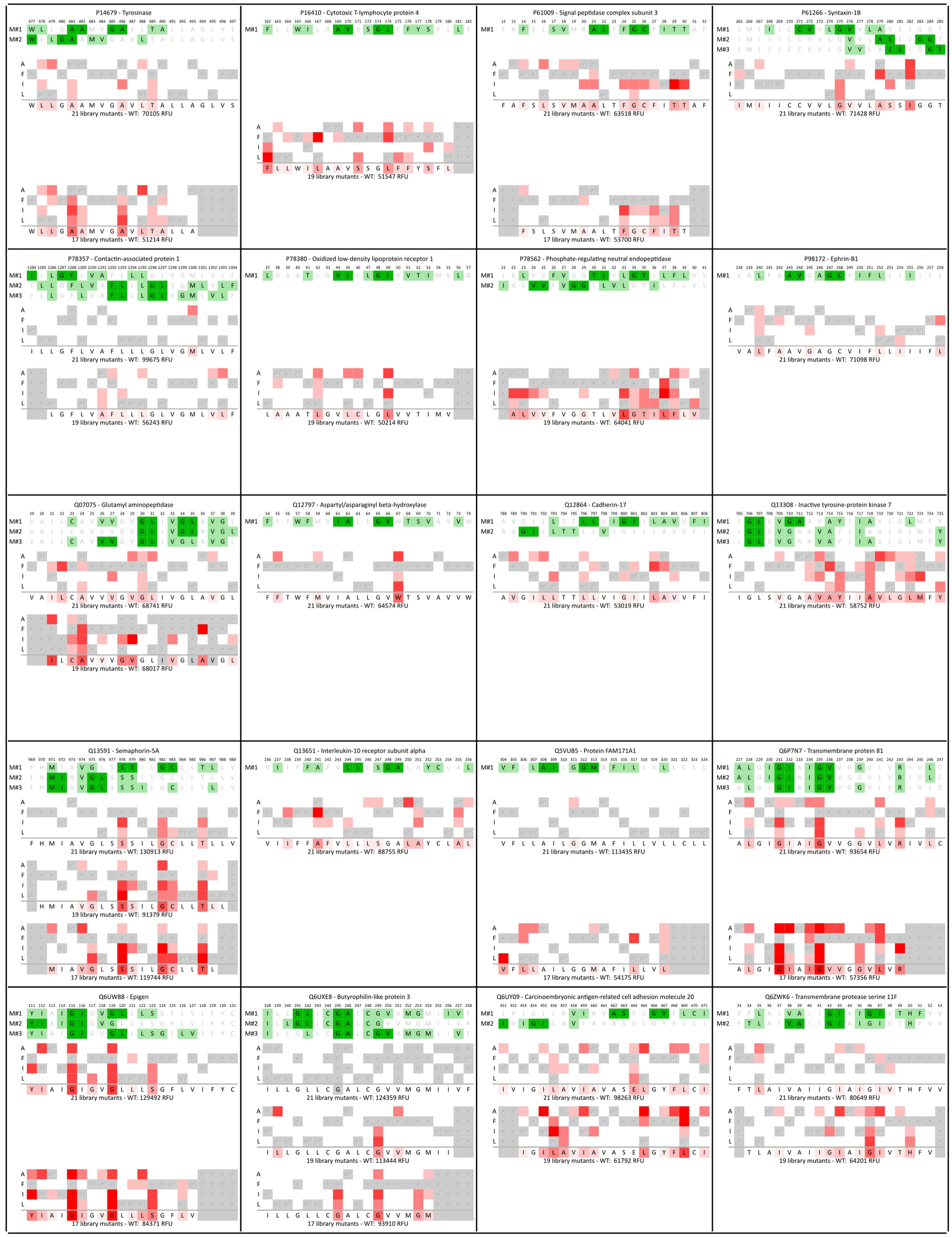
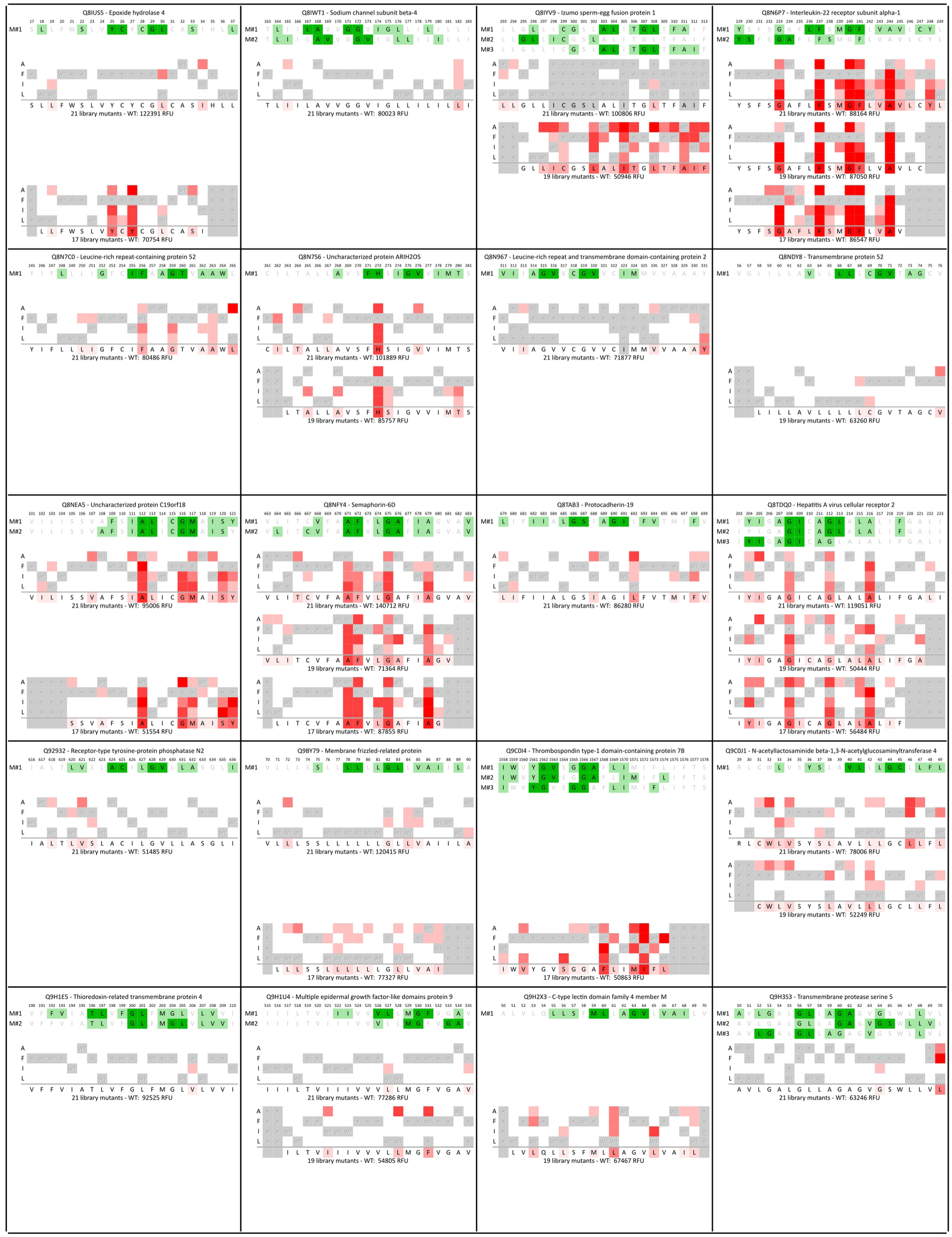
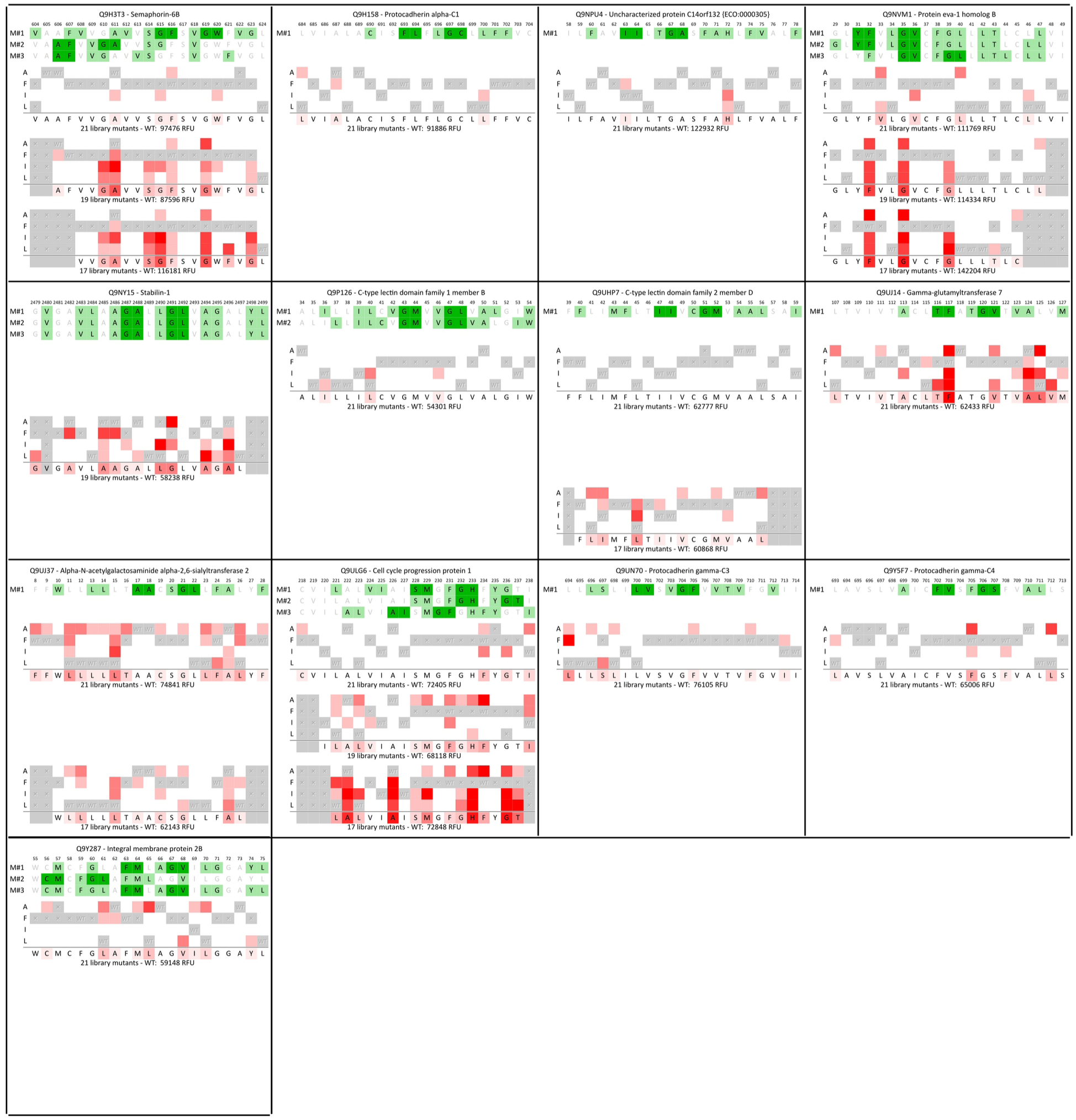
Mutational profiles of the predicted GAS_right_ constructs compared with the pattern of interfacial position predicted by CATM top three structural models. The mutational profiles were not computed for any constructs and lengths if the wild-type reconstructed fluorescence fell under a minimum threshold of 50,000 RFU.

**Table S3.** Summary of the classes of mutagenesis profiles. The number of predictions that are similar between CATM and AlphaFold3 are also reported.

| Mutagenesis result | Total | AF3 and CATM match<sup>1</sup> |  |
| --- | --- | --- | --- |
| Disruption matches CATM predicted interface | 12 | 9 | (75%) |
| Disruption maps to an alternative interface | 13 | 0 | (0%) |
| Disruption does not map to a clear interface | 6 | 1 | (17%) |
| Single polar disruptive position | 3 | 0 | (0%) |
| No or minimal disruption observed | 31 | 1 | (3%) |
| TOXGREEN signal too low for mutagenesis | 35 | 1 | (3%) |

<sup>1</sup>AF3 and CATM models are considered matching if their RMSD is under 2.2 Å

**Fig. S5.**
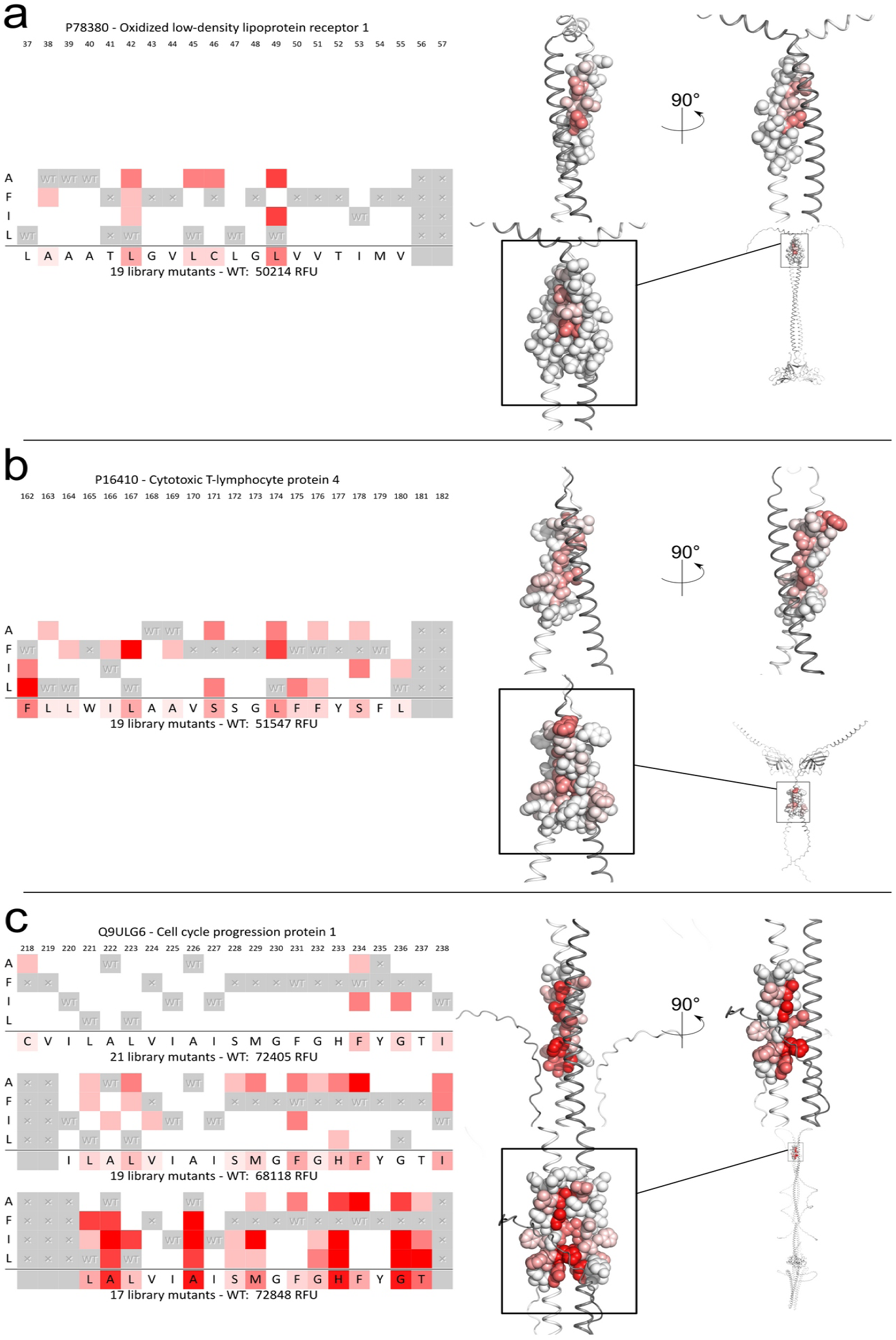
Mutagenesis profiles that match with an AlphaFold3 model. The three mutational profiles did not match the GAS_right_ configuration predicted by CATM but the interface highlighted by the mutagenesis is compatible with the dimeric AF3 structure.

**Fig. S5.**
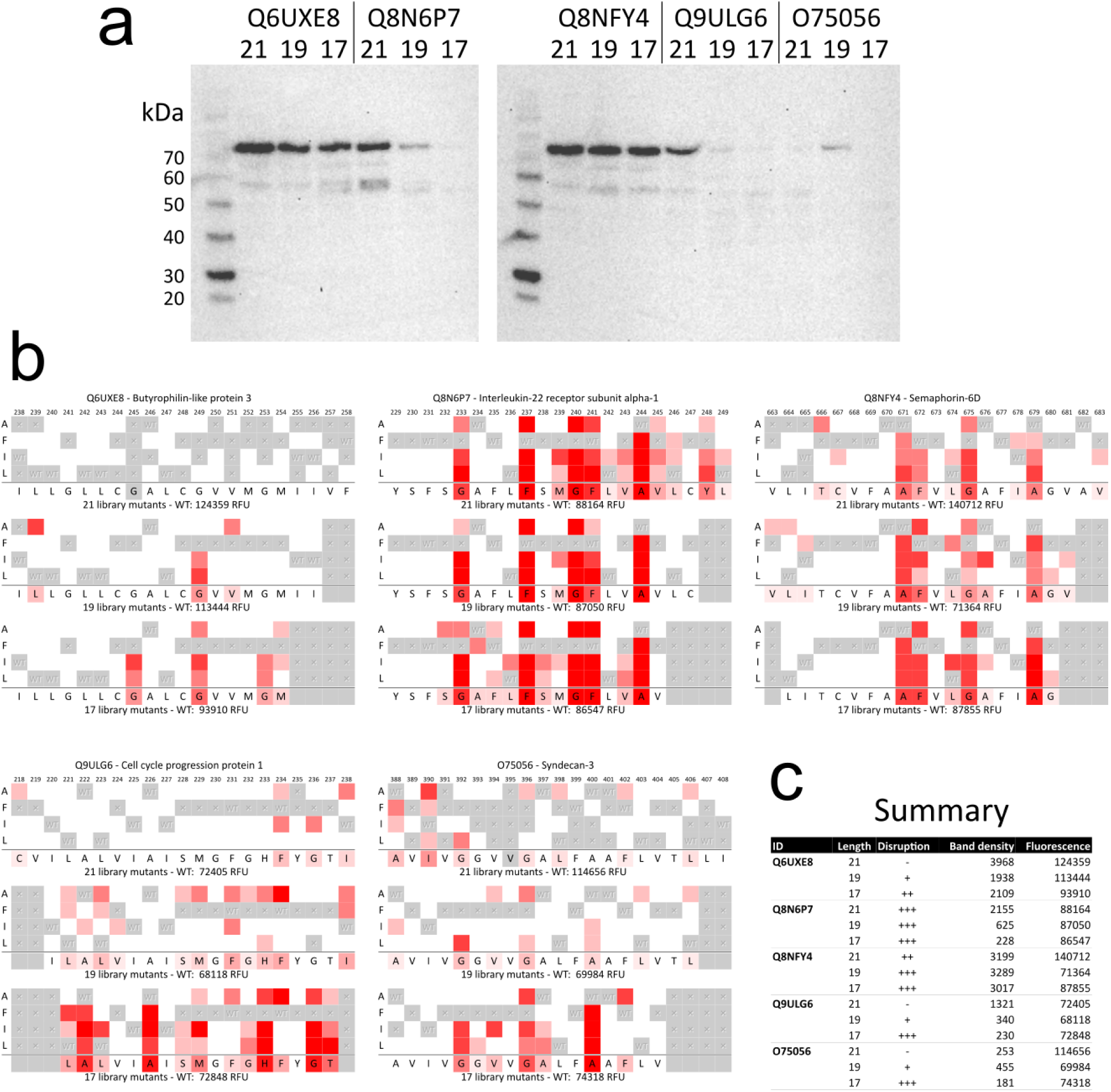
Western blot analysis does not reveal a trend between TOXGREEN signal, propensity to disruption and protein expression level. a) Western blots of all three lengths of five different TOXGREEN constructs using anti-maltose binding protein antibodies. b) Mutagenesis profiles of the same five constructs. c) Summary of TOXGREEN fluorescence, disruption level, and band density quantitation of the constructs.

**Table S4.** Series of steps that led to the selection of 100 predicted GAS_right_ constructs for the mutational analysis.

| Step | Constructs left |
| --- | --- |
| Human Single Pass Proteins Annotated in Uniprot | 2,383 |
| Potential GAS<sub>right</sub> structures predicted by CATM | 1141 |
| TM sequences that do not contain proline residues | 609 |
| TM sequences with annotated sequence length of 21 amino acids | 454 |
| CATM energy score below -10 kcal/mol | 363 |
| Structural models without significant interactions in juxtamembrane region | 122 |
| Sequences with not more than one polar residue | 110 |
| Sequences selected for experimental analysis (top 100 energy scorer) | 100 |

**Fig. S7.**
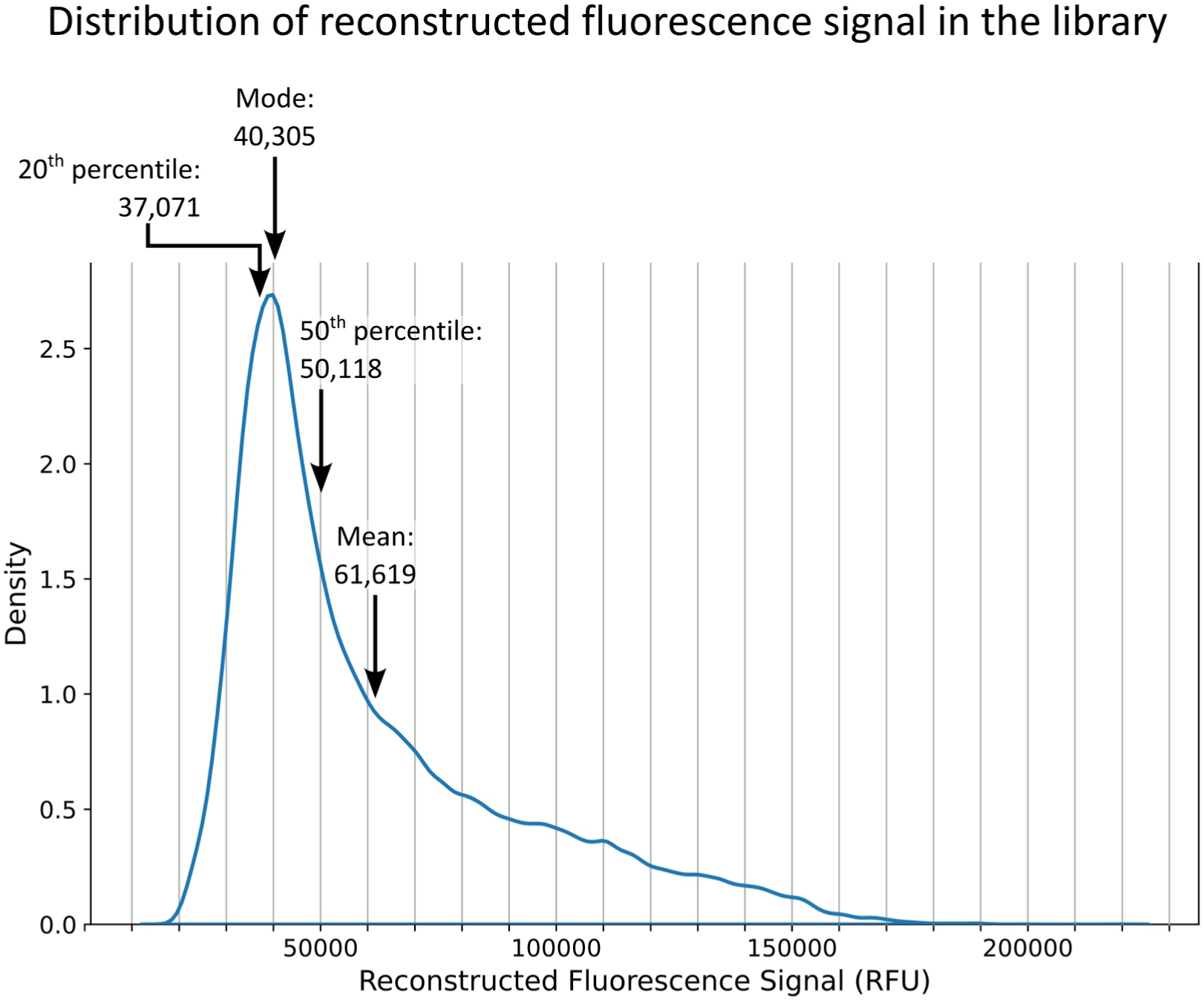
Distribution of reconstructed fluorescence in all constructs of the library. The distribution informed the choices for thresholds for the mutational analysis. A minimum threshold of 50,000 RFU (50^th^ percentile) was set for performing the mutagenesis analysis to guarantee sufficient dynamic range. A threshold of 35,000 RFU (∼ 20 ^th^ percentile) was chosen as the monomeric baseline threshold for the classification in the mutational analysis (see Methods).

